# A 3D atlas of sexually dimorphic lumbosacral motor neurons that control and integrate pelvic visceral and somatic functions in rats

**DOI:** 10.1101/2024.04.16.589836

**Authors:** John-Paul Fuller-Jackson, Ziying Yang, Nicole M Wiedmann, Alan Watson, Nathaniel EC Jenkins, Janet R Keast, Peregrine B Osborne

## Abstract

The lumbosacral spinal motor system comprises somatic motor neurons (SMNs) that target striated muscle and visceral motor neurons (VMNs; autonomic preganglionic neurons) that target peripheral ganglia. The brain coordinates these pathways to mediate functions such as continence, voiding and sexual function by ensuring appropriate changes of organ state with striated muscle activity (e.g., sphincter opening, relevant postural changes). These spinal circuits are important therapeutic targets for bioelectronic medicine yet are poorly defined in comparison with limb SMNs. We aimed to define the structural features and relationships between SMNs and VMNs relevant to pelvic function in male and female rats, by building 3D maps of lumbosacral spinal cord. We achieved this by applying large-volume immunostaining (choline acetyltransferase), with tissue clearing and advanced whole mount microscopy (light sheet and ribbon scanning confocal microscopy). We identified VMNs specific to pelvic organ function by microinjecting cholera toxin (beta-subunit) into the major pelvic ganglia (MPG). These VMNS are primarily located in segments L1-L2 (dorsal commissural nucleus) and L6-S1 (intermediolateral nucleus). Unexpectedly, small groups of SMNs in the urethral rhabdosphincter, cremaster and levator ani motor pools also projected through the MPG. Motor neuron counts and analysis of dendritic fields detected sexual dimorphism in both SMNs and VMNs. Their colocation of dendritic bundles suggested a mechanism of coordinating activity. This study has provided the first multiscale 3D atlas of rat lumbosacral cord. This will be shared as a resource on an open science platform (sparc.science) to allow further exploration and modeling of network features and cellular morphology.

## Introduction

The spinal cord motor neuron (MN) system is intensively studied as a model of the development and maturation of neural circuits (Blum and Gitler, 2022; Dasen, 2022). It is formed by two classes of motor neurons (MNs): *somatic* motor neurons (SMNs) that target striated muscle, and *visceral* motor neurons (VMNs) that target peripheral ganglia. VMNs are the preganglionic neurons in the autonomic nervous system, and input onto peripheral autonomic ganglion MNs that control the visceral organs, cardiovascular system, and other targets that are not striated muscle. The adult spinal motor system is hierarchically organized into longitudinal columns of neurons, with SMNs extending across the neuraxis of the spinal cord and VMNs forming non-contiguous columns interrupted by the cervical and lumbar enlargements (Barber et al., 1984). Each column controls a major body segment via pools of MNs that output to a common functional unit. This organization gives rise to spinal cord motor subsystems that control and coordinate patterned activity in functional groupings of striated muscles or autonomic motor targets.

The pelvic motor subsystem, located in the lumbosacral spinal cord, controls the pelvic and urogenital regions during excretory, sexual, and reproductive behaviors (Holstege and Collewijn, 2009; Thor and de Groat, 2010; de Groat, 2018). It produces complex, stereotypic rhythmic motor patterns — such as micturition, defecation, and ejaculation — that require a high degree of cooperativity between the VMNs controlling visceral activity and the SMNs controlling specialized striated muscles in the sexually dimorphic pelvic floor and urogenital region. The outputs are driven by a series of spinal cord motor pattern generators under reflexogenic control by spinal and brainstem circuits (Steuer and Guertin, 2019). However, motor patterns such as voiding during micturition and scent marking are also subject to psychogenic (conscious) controls by brain circuits that integrate these outputs with social and other behaviors (Hou et al., 2016; Keller et al., 2018).

Neural tracing in rodents has identified two groups of pelvic VMNs (separated by the lumbar enlargement) that are retrogradely labelled from their primary output target, the bilateral major pelvic ganglia (MPG) (Hancock and Peveto, 1979a, 1979b). The rostral group, mostly located in the midline dorsal commissural nucleus (DCN) of segments L1-L2, are sympathetic VMNs that output to the MPG via the hypogastric nerve. The caudal group form the sacral intermediolateral nucleus (IML) in the L6-S1 transition area and are parasympathetic VMNs that output to the MPG via the pelvic nerve (Anderson et al., 2009). Neural tracing has also identified pools of pelvic SMNs targeting the highly specialized, atypical, striated muscles in the pelvic floor and perineal regions — for example, the urethral rhabdosphincter that constricts the urethra and controls urine flow (Schrøder, 1980; Thor and de Groat, 2010). Pelvic SMNs are somatotopically organized into motor pools located in the lumbosacral transition, but the male cremaster muscle SMN group is located rostrally in the L1-L2 segments (Kojima et al., 1983). Pelvic SMNs are found in all three major columns specified during development (lateral, medial and hypaxial) (Dasen, 2022) and some motor pools are split across columns (Schrøder, 1980).

Our current understanding of the 3D neuroanatomical organization of the pelvic motor subsystem is almost entirely derived from reconstructions of sectioned spinal cords. Recent technical advances have improved these workflows (Fiederling et al., 2021), but the length of the spinal cord is a challenge for producing complete reconstructions from ordered contiguous series of spinal cord sections. Here we address these limitations by using advanced whole mount immunofluorescence and 3D microscopy (Belle et al., 2017; Fuller-Jackson et al., 2021b; Blain et al., 2023) to reveal the complex macroscopic neuroanatomical organization of pelvic motor neurons in the lumbosacral spinal cord. We also compare the spinal cord in female and male rats as previous evidence of extensive morphological sex differences in the pelvic motor system have not been examined in 3D studies (Anderson et al., 2009; Forger, 2009; Thor and de Groat, 2010; Oti and Sakamoto, 2023).

## Materials and Methods

### Animals

This study used fourteen male and ten female young adult Sprague-Dawley rats (8 – 10 weeks of age, males 300 – 350 g, females 200 – 250 g). The estrus cycle stage of female rats was not recorded. All animal procedures were approved by the Animal Ethics Committee of the University of Melbourne and complied with the Australian Code for the Care and Use of Animals for Scientific Purposes (National Health and Medical Research Council of Australia). Rats of the same sex were housed in groups of two or more under a 12/12 h light/dark cycle with ad libitum food and water.

### Injection of neural tracer into major pelvic ganglia (MPG)

The paired MPGs are located on the surface of the dorsolateral lobe of the prostate (male) or uterine cervix (female). Spinal cord neurons projecting to the MPG were identified by retrograde neural tracing with cholera toxin subunit B (CTB). Six rats (three male and three female) were anesthetized with isoflurane (3% in oxygen for induction, 1–2% for maintenance), followed by a midline ventral incision to access the MPGs. Each MPG was separated from underlying tissue by blunt dissection with fine angled forceps, and a sterile 2 mm square piece of parafilm inserted underneath to minimize leakage of tracer. A solution of CTB (low-salt, 0.3% w/v; List Biological Labs, CA, USA) and Evans Blue (0.05% w/v; Sigma-Aldrich, NSW, Australia) in sterile water was then microinjected into the MPG using a glass pipette attached to a Picospritzer (Parker Hannifin). The total volume injected across three microinjections was 1.8 – 2.2 μl. After injection, the pipette was held in place for 5 seconds, and then withdrawn and the site washed with sterile saline.

Following microinjection of both MPGs, the abdominal wall was sutured closed, and the skin closed with surgical staples. Postoperative analgesia included buprenorphine (Clifford Hallam Healthcare, VIC, Australia; 0.05 mg/kg, at the time of surgery and 8 h postsurgery) and meloxicam (Troy Laboratories, NSW, Australia; 1 mg/kg, 8 and 24 h postsurgery), administered subcutaneously. Post-surgery no adverse events were observed. Three days after surgery tissues were collected.

### Tissue collection

Rats were anesthetized (ketamine 100 mg/kg and xylazine 10 mg/kg, i.p.) prior to tissue fixation via intracardiac infusion, as detailed previously (Keast et al., 2020). Briefly, rats were perfused with 0.9% saline containing 1% sodium nitrite and 5000 IU/ml heparin for 3 min, then 4% paraformaldehyde in 0.1 M phosphate buffer (pH 7.4) for 10 min. The spinal cord was removed and postfixed for 1 h, followed by three 1 h washes in 0.1 M phosphate buffered saline (PBS, pH 7.2). Tissue was stored in PBS with 0.1% sodium azide at 4°C. In one female rat from the CTB injection study, the upper lumbar cord was damaged at dissection so was not analyzed further.

### Immunolabelling and tissue clearing of whole mount spinal cord

Fixed spinal cords from 4 female and 7 male rats were sub-dissected to isolate thoracolumbar segments T13 to L3 and lumbosacral segments L4 to S3. A longer sample of spinal cord (spinal segments T3 to S3) was also prepared from one male rat. Segment boundaries were identified by determining the position of the most caudal of the emerging rootlets of each ventral root (Watson and Kayalioglu, 2009; Watson et al., 2009). This fiduciary marker could also be identified by light sheet microscopy from the raw signal and was used to identify segments in 3D spinal cord images.

Large-volume clearing and immunolabelling was performed as previously described (Fuller-Jackson et al., 2021b), using an iDISCO based workflow (Renier et al., 2014; Belle et al., 2017). Spinal cords were washed in 1x Dulbecco’s PBS (DPBS; Sigma-Aldrich, NSW, Australia; 6 x 15 min), dehydrated in progressively increasing concentrations of methanol (50%, 80% and 100% in DPBS; 1.5 h each), incubated overnight in 6% hydrogen peroxide in methanol at 4°C, then rehydrated in decreasing concentrations of methanol (100%, 100%, 80%, and 50%; 1.5 h each). After incubation in DPBS (1.5 h), spinal cords were transferred to a blocking solution for 36 h at room temperature (DPBSG-T: DPBS with 0.2% gelatin, 0.5% Triton X-100, and 0.01% thimerosal). Spinal cords were incubated with primary antibodies in DPBSG-T with an additional 0.1% saponin for 10 d at 37°C, washed (DPBS-T; DPBS with 0.5% Triton X-100; 6 x 15 min), followed by incubation with secondary antibodies in DPBSG-T with 0.1% saponin for 4 d at 37°C. Continuing at room temperature, spinal cords were rinsed in DPSB-T (6 x 15 min), followed by dehydration in methanol in DPBS (20%, 40%, 60%, 80%, 2 × 100%; 1 h), incubation in 66% dichloromethane and 33% methanol overnight. Finally, spinal cords were washed three times with 100% dichloromethane for 30 min, optically cleared with dibenzyl ether, then stored in fresh dibenzyl ether.

Details of primary and secondary antibodies are provided in Table 1. All spinal cords were immunolabelled for choline acetyltransferase (ChAT), with the exception of one spinal cord that was immunolabelled with the neuronal nuclear antigen, NeuN (also known as Fox-3). Spinal cords from the CTB microinjection study were immunolabelled for CTB and ChAT.

**Table 1:**
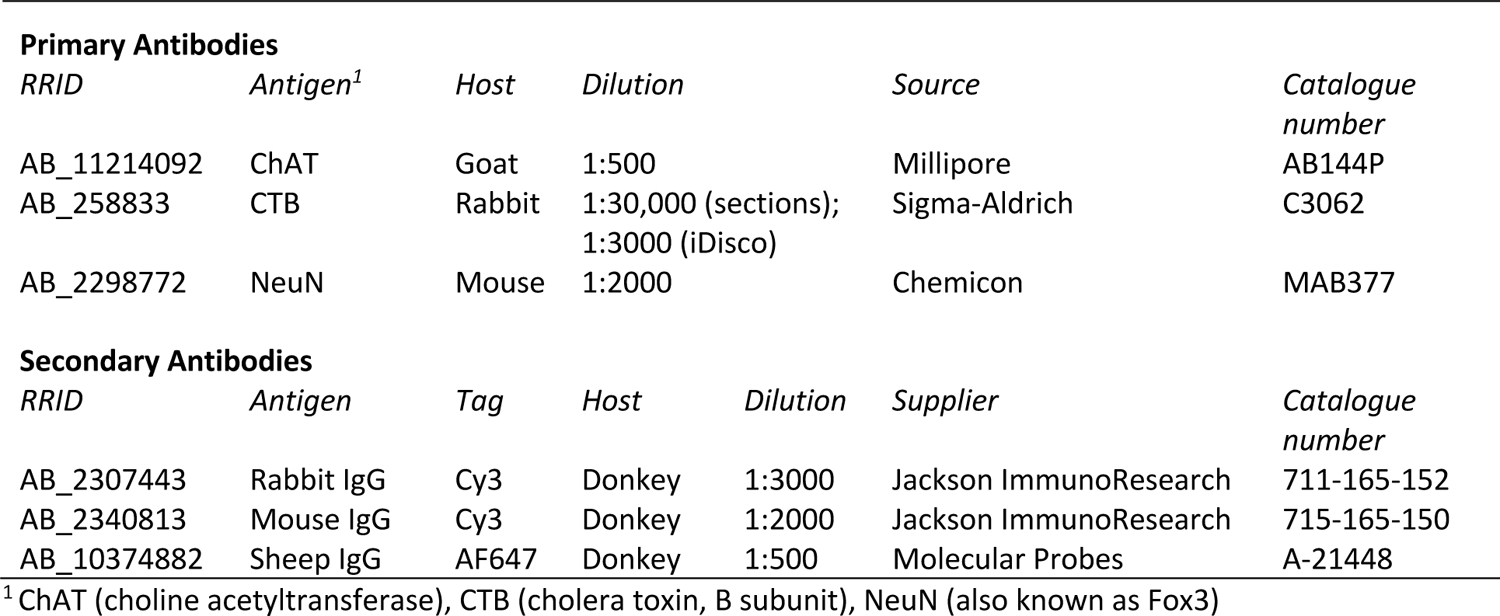
Antibodies used for immunofluorescence microscopy.

### Large volume 3D microscopy

Cleared spinal cords were transferred to ethyl cinnamate and visualized via light sheet microscopy (Ultramicroscope II, Miltenyi Biotec, Germany), using either a zoomable 2x lens (MVPLAPO, Olympus, Japan) or a 12x fixed zoom lens (LVMI PLAN, LaVision, MI, USA). Single-sided 3-sheet illumination was used with lasers 561 and 639 nm paired with emission filters 620/60 and 680/30 nm. The numerical aperture of the light sheet was set to 0.156. Images were acquired with 2 μm z-steps and 50-200 ms exposure. The image acquisition area was reduced by ∼25% to minimize the impact of the light sheet waist on the uniformity of the image during mosaic acquisitions, which were performed with 10% overlap between image stacks. Image stacks were converted to Imaris file types (Imaris File Converter, Bitplane) and stitched using Imaris Stitcher, prior to detailed visualization in Imaris (RRID:SCR_007370). 3D representation of motor nuclei was achieved using the *Surfaces* function in Imaris, with contours drawn manually around motor nuclei in either transverse, sagittal or horizontal optical slices.

Two cleared spinal cords labelled with ChAT were visualized via ribbon scanning confocal microscopy (RSCM; Caliber, MA, USA) as previously described (Watson et al., 2017). The microscope was fitted with a Nikon CFI90 20x glycerol-immersion objective (Nikon, NY, USA) with 8.3 mm working distance.

Volumes were captured with voxel resolution of 0.491 × 0.491 × 10.67 μm (x, y, z). Laser intensity and detector settings were specific to each sample based on the levels of staining. In all cases, the intensity of the laser was increased by linear interpolation throughout deeper focal planes to compensate for absorption of excitation and emission light. Images acquired in this way were stitched and assembled into composites using a 24 node, 608 core cluster, then converted into the Imaris file format. Volumes were rendered using Imaris.

### Quantitation of spinal neurons in specific nuclei

VMNs and SMNs were manually counted in raw images from cleared spinal cord whole mounts by using Neurolucida 360 (MBF Bioscience, RRID:SCR_016788) in the 2D view to mark single neurons that were in regions identified in horizontal 2 μm thick virtual slices. Pelvic VMNs were identified by appearance of CTB in their somata as a result of retrograde neural tracer injections into the MPGs. In L6-S1 segments, ChAT^+^/CTB^+^ and ChAT^+^/CTB^-^ neurons were counted in the IML on one side of the cord, and the ratio of counts used to calculate efficiency of retrograde labeling. In the L1-L2 segments, the neuron classes were counted in the principal IML on one side and in the midline DCN, but retrograde labeling efficiency was not estimated, as these regions do not exclusively target the MPG and contain other classes of non-pelvic VMNs. We also identified CTB^+^ SMNs in some regions (Cr9, ExU9 and Tail9 nuclei), which were counted manually.

### Morphological analysis of anal and urethral rhabdosphincter motor nuclei

Urethral and anal rhabdosphincter-innervating motor nuclei were analyzed in six rats of each sex. Unlike humans and non-human primates, in this species these two functional classes are each located in a distinct nucleus within the lumbosacral cord (Nadelhaft and McKenna, 1987). These sections were immunolabelled for a previous study (Fuller-Jackson et al., 2021b), however in the current study this open dataset was reanalyzed to quantify the area of each motor nucleus in each section. Lumbosacral spinal cords (L5-S2) were cryoprotected overnight in 0.1 M PBS containing 30% sucrose prior to embedding in Tissue-Tek optimal cutting temperature compound (Sakura Finetek, CA, USA). Transverse cryosections (40 μm) of the entire block were collected in order, with alternate sections immunolabelled for ChAT. Free-floating sections were washed in 0.1 M PBS, blocked in 0.1 M PBS with 10% normal horse serum (NHS) and 0.5% Triton X-100 for 2 h, washed in 0.1 M PBS, then incubated in solution containing primary antibodies, 2% NHS, 0.5% Triton X-100 and 0.1% sodium azide for 48 h at room temperature.

After washing in 0.1 M PBS, sections were incubated in species-specific secondary antibodies tagged with AF647 in a solution of 0.1 M PBS containing 2% NHS, and 0.5% Triton X-100 for 4 h at room temperature. Mounted on glass-slides in rostrocaudal anatomic order, sections were cover-slipped using carbonate-buffered glycerol (pH 8.6). Primary and secondary antibodies are described in Table 1.

All spinal cord sections were imaged (10x objective, pixel scaling 0.645 x 0.645 μm, five z-steps at 5 μm) with a Zeiss AxioImager Z1 (Carl Zeiss Microscopy). Sections were aligned in TissueMaker (MBF Bioscience, PA, USA; RRID:SCR_017322), producing a reconstructed spinal cord dataset (Fuller-Jackson et al., 2021a). The distance between sections was 80 μm. In Neurolucida 360, the urethral and anal rhabdosphincter-innervating motor nuclei (ExU9, ExA9) were identified, and the outside of each nucleus manually traced in each section using the *Contour* function. This involved tracing around the collection of neurons as they appear in the section and was performed bilaterally in each section containing these nuclei. For ExA9, the entire rostrocaudal extent of this nucleus was mapped. For ExU9, only sections where ExU9 was the sole nucleus in the ventrolateral region were mapped. This excluded rostral sections that contained both ExU9 and Gl9/Hm9 in the same region, which were not mapped due to the difficulty in accurately distinguishing the boundaries of each nucleus. In Neurolucida Explorer (MBF Bioscience; RRID:SCR_017348), the area of each contour on each section was determined across all relevant sections. Data were plotted as motor nucleus area (μm^2^) from the most rostral section to the most caudal section. In each animal, the total contour area from each nucleus was then calculated and compared between male and female groups.

### Analysis of dendritic fields of caudal lumbosacral visceral (preganglionic) MNs

The distribution of CTB in parasympathetic preganglionic neurons in L6-S1 segments was used to visualize the features of their dendritic fields in 3D in cleared spinal cord of male and female rats. In dendrites, CTB granules were punctate in appearance. Using the *Spots* function in Imaris, spherical 3D objects were detected according to their size (1 μm XY, 2 μm Z) and signal intensity after background subtraction. Due to the punctate nature of CTB within neuronal processes, individual dendrites could not be traced.

To generate the total volume of these dendritic fields, CTB objects in Imaris were filtered prior to the fitting of a convex hull around the remaining objects. Alpha Shapes creates polygons from objects according to a set parameter (α). The higher the α, the greater the distance between objects that can be included in the creation of polygons using *Alpha Shapes*. For this study α was set to 3-4, based on visual assessment of each dataset. This effectively excluded distant objects that were isolated from other objects. The final step, the fitting of a convex hull, involved the generation of multiple smaller convex hulls along the length of the dataset. Initially a single convex hull was applied, however each of the most distant CTB points in any given direction created a convex hull that overestimated the volume of the dendritic field. Smaller convex hulls for subsets of CTB points sorted by Z were generated, with 20-30% overlap of CTB points between each adjacent convex hull. The union of the smaller convex hulls improved the mapping of the contours of dendritic projections. The overlaps between convex hulls were removed and these convex hulls union fused to produce a single watertight volume.

Our method was developed as a Python pipeline which is assembled by open-sourced Python libraries and our own algorithms (https://gitlab.unimelb.edu.au/lab-keast-osborne-release/3d-points-volume-estimation). The Python library, PyMeshlab (Muntoni and Cignoni, 2021), was employed to create Alpha Shape in the initial step. The filtered spots were then used to compute all the described smaller Convex Hull using Trimesh (Dawson-Haggerty et al., 2019). Finally, the union of the Convex Hull shapes were processed by PyMeshlab. We also used Numpy (Harris et al., 2020) and Pandas (The pandas development team, 2020) in our program.

### Tracing axon projections from sacral SMNs

Axons of sacral (S1) SMNs were traced within the spinal cord white matter. These included axons that projected laterally into the white matter tracts, before turning perpendicularly to continue either rostrally or caudally to the next segment. Cleared lumbosacral spinal cord immunolabelled with ChAT was imaged via light sheet microscopy at 12x magnification and the resulting Imaris files converted to JPEG2000 using Microfile+ (MBF Bioscience, RRID:SCR_018724). Using Neurolucida 360’s *Tree tracing* function in the 3D environment, axons of sacral SMNs were traced, by performing user-guided tracing with the *Directional Kernels* algorithm of choice. Where two traced axons were in close proximity, the Smart-manual tracing tool was used, thereafter returning to user-guided tracing. Axons were traced as far as they could be confidently identified as the same axon.

### Antibody characterization

Primary antibodies were obtained from the following sources (information on specificity was obtained from the manufacturer): ChAT (Millipore; AB144P; batch 2971003; RRID: AB_11214092): immunoaffinity purified polyclonal antibody raised in goat against the human placental enzyme; In Western blotting, 70-kDa bands were detected in NIH/3T3 lysate, matching the molecular weight of ChAT protein.

NeuN (Chemicon; MAB377; batch 2453249; RRID: AB_2298772): purified monoclonal antibody raised in mouse against purified cell nuclei from mouse brain. In Western blotting, recognized 2-3 bands in the 46-48 kDa range, matching molecular weight of NeuN.

CTB (Sigma-Aldrich; C3062; batch 048M4780V; RRID: AB_258833): polyclonal antibody raised in rabbit against Vibrio cholerae, reacting with cholera toxin but not staphylococcal enterotoxin A, staphylococcal enterotoxin B, and pseudomonas exotoxin A.

### Experimental design and statistical analysis

For the qualitative description and representation of spinal cord motor nuclei, long (T3-S3) and short (L4-S3) spinal cords from male rats were used. One male and one female rat spinal cord (L5-S2) were used to demonstrate sexually dimorphic motor nuclei in the intact cord, while cryosections from spinal cords of six male and six female rats were used for the quantitative comparison of these motor nuclei. Plots of the total area of motor nuclei contours in sections across the neuraxis of L6 were used to compare the size of the motor nuclei in male and female rats. Spinal cords from three male and three female rats that received MPG microinjections of CTB were used for the quantitative comparison of neuron number and dendritic field volume of sacral preganglionic neurons. Comparisons between sexes were made using a two-tailed *t-*test. Graphs were generated in JMP 16.0 (SAS Institute, NC, USA; RRID:SCR_014242)) and data presented as group mean and individual subjects. Statistical analyses were performed using JMP 16.0.

### Figure production

For representative images of light sheet datasets, neurons were visualized with γ set in Imaris to 1.4 (ChAT, CTB) or 1.6 (NeuN). To best demonstrate neuronal morphology in 2D outputs from Imaris, small linear adjustments were made to levels using Photoshop (Adobe Creative Suite v2023). In some figures, the ventral roots have been non-destructively removed from view using Photoshop, particularly when intense ChAT staining of roots distracts from fine neuronal features within the spinal cord image. Figures were constructed in InDesign (Adobe Creative Suite v2023).

### Data sharing

Raw data from this study will be published under an open access license on https://sparc.science/. Movies illustrating the complete 3D datasets are also available online (DOI for each Movie provided where relevant in the Results).

## Results

### Visceral (preganglionic) and somatic motor neurons (VMN, SMNs) in caudal spinal cord

To provide an initial overview of motor columns in caudal spinal cord, we used RCSM to identify ChAT^+^ neurons in large intact contiguous lengths of spinal cord that extended from segment S2 as far rostrally as segment T3 (Fig. 1A, Media 1 [https://doi.org/10.26188/25567014.v1). One male spinal cord was imaged at an XY resolution of 0.491 µm in optical slices spaced 10.7 µm apart in the Z axis. At this resolution, individual ChAT^+^ neurons could be easily defined and used as a chemoarchitectural marker to identify VMNs (preganglionic) and SMNs from segments T10 to S2 (Fig. 1B). Segment boundaries were identified by the caudal limit of rootlets within each ventral root (visible from background fluorescence in raw signal), which is the fiduciary marker most used by spinal cord atlases of rodents and other species (Watson and Kayalioglu, 2009; Watson et al., 2009). ChAT is not an exclusive marker of MNs as cholinergic interneurons are also located in intermediate laminae as well as the dorsal horn. However, the scattered isolated ChAT^+^ interneurons could be readily distinguished from MNs, as the latter were larger multipolar neurons that aggregated into columns or groups of neurons extending longitudinally through the intermediate grey matter or ventral horn (Fig. 1B). To show the 3D topology of the columns, we processed the raw signal with Imaris software and annotated images with a mesh surrounding each major MN column (Fig. 1B). This showed how the lateral motor columns extend through most of the lumbar enlargement, spanning the discontinuity in the intermediolateral preganglionic columns, i.e., between caudal thoracic (sympathetic) and caudal lumbar (parasympathetic) segments (Fig. 1B). This 3D approach also effectively demonstrated the complexity in rostrocaudal organization of individual subcolumns of MNs (Schrøder, 1980; Barber et al., 1984), especially between segments L5-S1, where numerous aggregates of SMNs partially overlap in their segmental distribution and alignment with VMNs. The cellular features of these different visceral and somatic MN classes are demonstrated in Figs. 1C-E.

**Figure 1.**
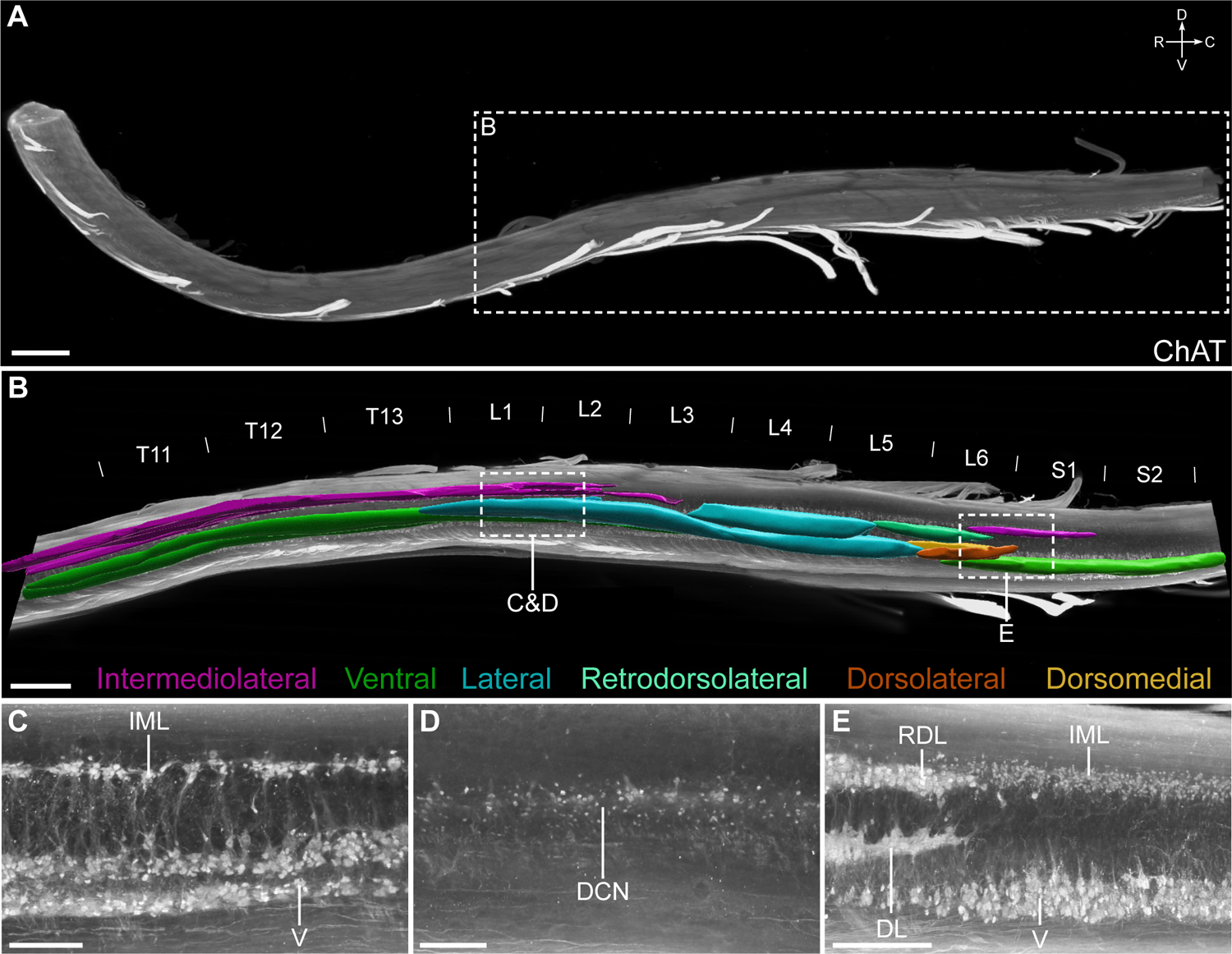
Distribution of motor columns and pools in the caudal male rat spinal cord. ***A***, Intact, cleared spinal cord containing segments T3 to S2 immunolabelled with choline acetyltransferase (ChAT) and visualised with ribbon scanning confocal microscopy. ***B***, 3D segmentation of motor pools shown in a sagittal orthogonal projection of the caudal spinal cord. From the same dataset, example orthogonal projections in sagittal plane of upper lumbar cord showing ***C***, Intermediolateral nucleus (IML), ***D***, Dorsal commissural nucleus (DCN) and ***E***, Lumbosacral IML. The DCN was unable to be segmented due to its indistinct boundaries and no capacity to distinguish preganglionic neurons from ChAT^+^ interneurons. L, lateral; V, ventral; RDL, retrodorsolateral; DL, dorsolateral; DM, dorsomedial (Schrøder, 1980). Scale bars: A 2000 μm, B 1000 µm, C-E 300 μm.

### Retrograde tracing identifies pelvic VMNs and reveals somatic motor pathways projecting via the MPG

Spinal VMNs controlling pelvic organ function are split into two functionally distinct groups of sympathetic and parasympathetic preganglionic neurons, which in rat are respectively located either side of the lumbar enlargement in L1-L2 and L6-S1 cord. SMNs that innervate striated muscle related to urogenital or lower bowel function (urethral and anal sphincters, pelvic floor) are also located in the caudal lumbosacral segments, except for a smaller rostral group in segments L1-L2 that target cremaster muscle.

To visualize the 3D structure and location of both sympathetic and parasympathetic pelvic preganglionic neurons, we injected the MPG of 3 male and 3 female rats with the retrograde neural tracer CTB (Fig. 2A). In rodents, this ganglion contains most of the autonomic neurons that innervate tissues of pelvic organs and is the site where both sympathetic and parasympathetic preganglionic neurons synapse. These preganglionic axons project via the hypogastric and pelvic nerves (Fig. 2A). Spinal neurons labelled by CTB injection into the MPG are also shown in Media 2 (https://doi.org/10.26188/25479250.v1) and Media 3 (https://doi.org/10.26188/25484116.v1). In the upper lumbar region of spinal cord, CTB^+^/ChAT^+^ preganglionic neurons were restricted to neuronal columns in segments L1-L2 (Fig. 2B, Media 3 [https://doi.org/10.26188/25484116.v1]) and were more densely distributed in the midline DCN than in the lateral IML. In the lumbosacral IML (L6-S1), the source of parasympathetic pelvic preganglionic neurons, most of these ChAT^+^ neurons were labelled by CTB (Fig. 2B, Media 2 [https://doi.org/10.26188/25479250.v1]), indicating the effectiveness of our retrograde tracing method. A minority of SMNs in specific locations were also labelled (discussed below).

**Figure 2.**
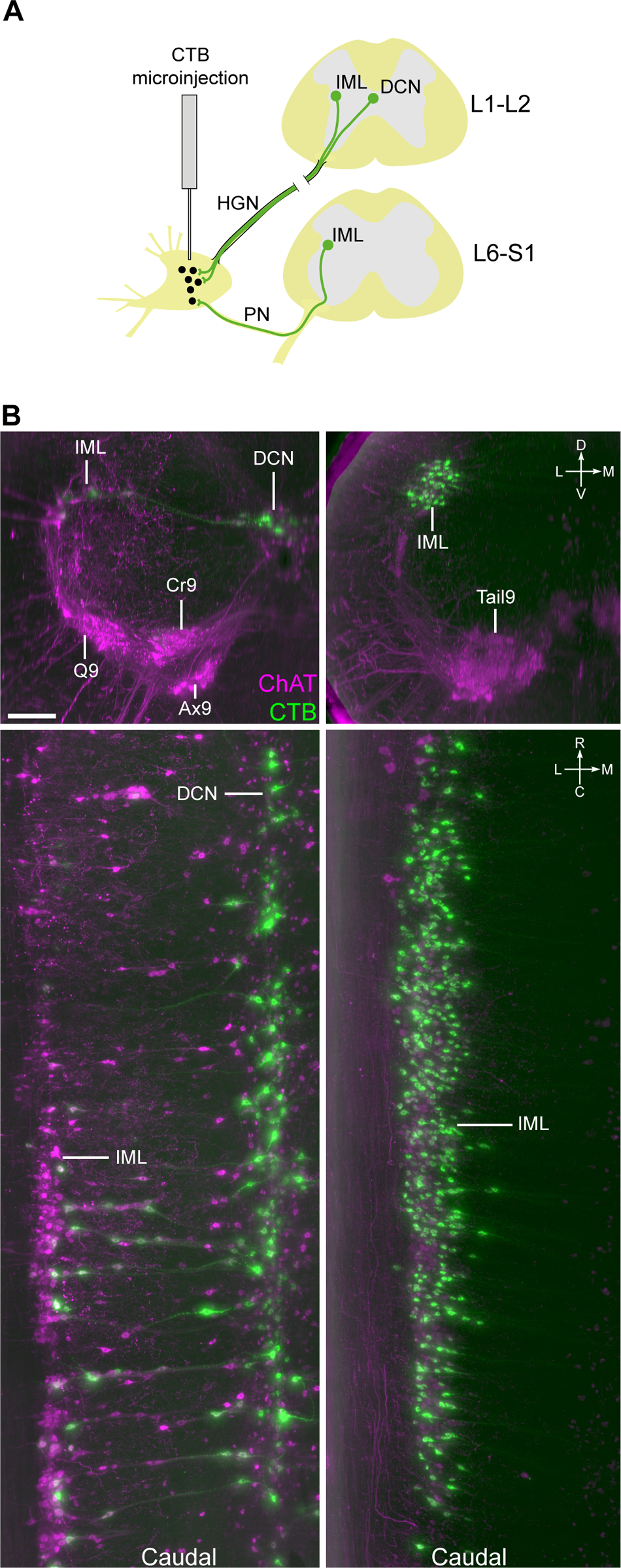
Retrograde labelling of preganglionic neurons in the lumbosacral spinal cord that project to the major pelvic ganglia. ***A***, Schematic representation of cholera toxin subunit B (CTB) microinjection into the rat major pelvic ganglion with resulting retrograde labelling in preganglionic neuron populations in spinal cord. ***B***, Virtual slices of CTB-filled preganglionic neurons in cleared L1-L2 (left) and L6-S1 (right) spinal cord (male). Spinal cord immunolabelled for CTB and choline acetyltransferase (ChAT). Top panels show spinal cord in transverse orientation, whereas lower panels show the columns of preganglionic neurons in the rostrocaudal axis. Images oriented with midline to the right. HGN, hypogastric nerve; PN, pelvic nerve; IML, intermediolateral nucleus; DCN, dorsal commissural nucleus; Q9, quadriceps; Cr9, cremaster; Ax9, axial; Tail9, tail. Scale bar: 200 μm (applies to all panels).

We then manually counted CTB^+^ neurons in preganglionic nuclei, analyzing the IML on one side and the entire DCN (Table 2*A*). In the L6-S1 IML (the sole location of parasympathetic preganglionic neurons), we also counted the entire population of ChAT^+^ neurons to provide an estimate of our CTB labelling efficiency. This indicated that 85.0 ± 3.3% (mean ± SEM, n=6, range 71-93%) lumbosacral preganglionic neurons were CTB^+^. Our analysis of the total preganglionic neuron population on one side of L6-S1 revealed a sex difference (male > females; males, 844 ± 30, n = 3 versus females, 707 ± 31, n =3; two-tailed *t-*test*, P* = 0.033)

**Table 2.**
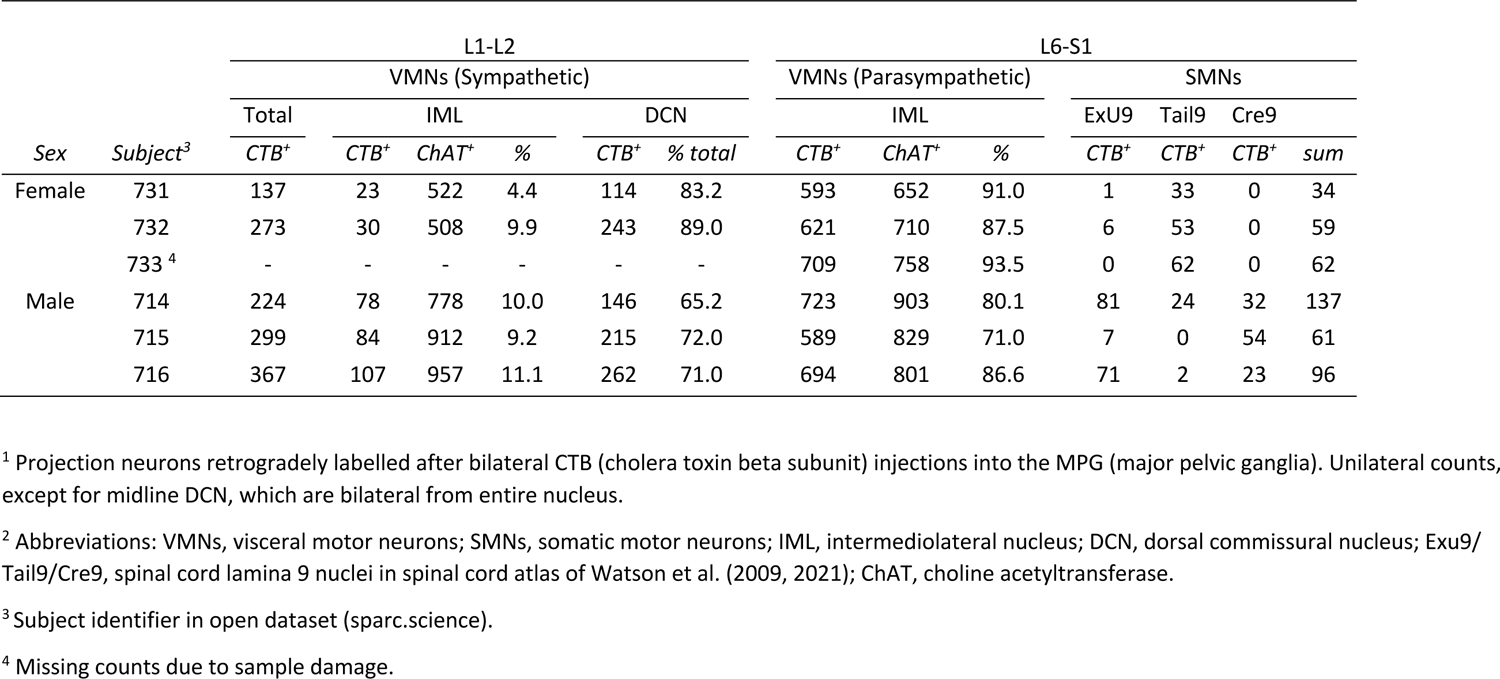
Counts of lumbosacral spinal cord VMNs (autonomic preganglionic MNs) and SMNs with projections to the major pelvic ganglia (MPG) ^1,2^.

| Sex | Subject<sup>3</sup> | L1-L2 |  |  |  |  |  | L6-S1 |  |  |  |  |  |  |
| --- | --- | --- | --- | --- | --- | --- | --- | --- | --- | --- | --- | --- | --- | --- |
|  |  | VMNs (Sympathetic) |  |  |  |  |  | VMNs (Parasympathetic) |  |  | SMNs |  |  |  |
|  |  | Total | IML |  |  | DCN |  | IML |  |  | ExU9 | Tail9 | Cre9 |  |
|  |  | CTB<sup>+</sup> | CTB<sup>+</sup> | ChAT<sup>+</sup> | % | CTB<sup>+</sup> | % total | CTB<sup>+</sup> | ChAT<sup>+</sup> | % | CTB<sup>+</sup> | CTB<sup>+</sup> | CTB<sup>+</sup> | sum |
| Female | 731 | 137 | 23 | 522 | 4.4 | 114 | 83.2 | 593 | 652 | 91.0 | 1 | 33 | 0 | 34 |
|  | 732 | 273 | 30 | 508 | 9.9 | 243 | 89.0 | 621 | 710 | 87.5 | 6 | 53 | 0 | 59 |
|  | 733 <sup>4</sup> | - | - | - | - | - | - | 709 | 758 | 93.5 | 0 | 62 | 0 | 62 |
| Male | 714 | 224 | 78 | 778 | 10.0 | 146 | 65.2 | 723 | 903 | 80.1 | 81 | 24 | 32 | 137 |
|  | 715 | 299 | 84 | 912 | 9.2 | 215 | 72.0 | 589 | 829 | 71.0 | 7 | 0 | 54 | 61 |
|  | 716 | 367 | 107 | 957 | 11.1 | 262 | 71.0 | 694 | 801 | 86.6 | 71 | 2 | 23 | 96 |

<sup>1</sup> Projection neurons retrogradely labelled after bilateral CTB (cholera toxin beta subunit) injections into the MPG (major pelvic ganglia). Unilateral counts, except for midline DCN, which are bilateral from entire nucleus.
<sup>2</sup> Abbreviations: VMNs, visceral motor neurons; SMNs, somatic motor neurons; IML, intermediolateral nucleus; DCN, dorsal commissural nucleus; Exu9/Tail9/Cre9, spinal cord lamina 9 nuclei in spinal cord atlas of Watson et al. (2009, 2021); ChAT, choline acetyltransferase.
<sup>3</sup> Subject identifier in open dataset (sparc.science).
<sup>4</sup> Missing counts due to sample damage.

Similar quantitation of CTB^+^ preganglionic neurons in the L1-L2 segments from the same animals (Table 2*A*) showed that fewer neurons were labelled with CTB than in the L6-S1 segments (L1-L2, 137-369; L6-S1, 589-723). The majority (65-89%) of the CTB^+^ preganglionic neurons in L1-L2 were in the DCN. In the L1-L2 IML, CTB^+^ neurons comprised only 4-11% of all preganglionic neurons identified by ChAT-immunoreactivity. Sex differences in these segments could not be analyzed due to the smaller number of biological replicates.

CTB injected into the MPG did not label ChAT^+^ interneurons or ChAT^-^ neurons in any spinal segments. However, unexpectedly, CTB labelled a subpopulation of ChAT^+^ SMNs in certain nuclei (Fig. 3A, Table 2*B*), which we deduce project to their peripheral targets via the MPG. These SMNs were identified using an atlas of the rat spinal cord (Watson et al., 2009, 2021). A comparison of 3 male rats and 2 female rats identified a sex difference in the distribution of CTB^+^ SMNs. In male rats, but not female rats, CTB labelled L1-L2 cremaster SMNs in Cr9 (Fig. 3B, D, Media 3 [https://doi.org/10.26188/25484116.v1]), the motor pool that targets the involuntary striated muscle that retracts the testicles, and L6 SMNs in ExU9 (Fig. 3C, E, Media 2 [https://doi.org/10.26188/25479250.v1] that contract the urethral rhabdosphincter. In both male and female rats, CTB also labelled S1 SMNs in Tail9 (Fig. 3B, F, Media 2 [https://doi.org/10.26188/25479250.v1]), which contains motor pools that target tail muscles and the levator ani muscle in the pelvic floor. All CTB/ChAT^+^ SMNs had large rounded somata with the multipolar morphology characteristic of alpha SMNs (Friese et al., 2009; Burke, 2016) and not the small spindle-shaped morphology of gamma SMNs.

**Figure 3.**
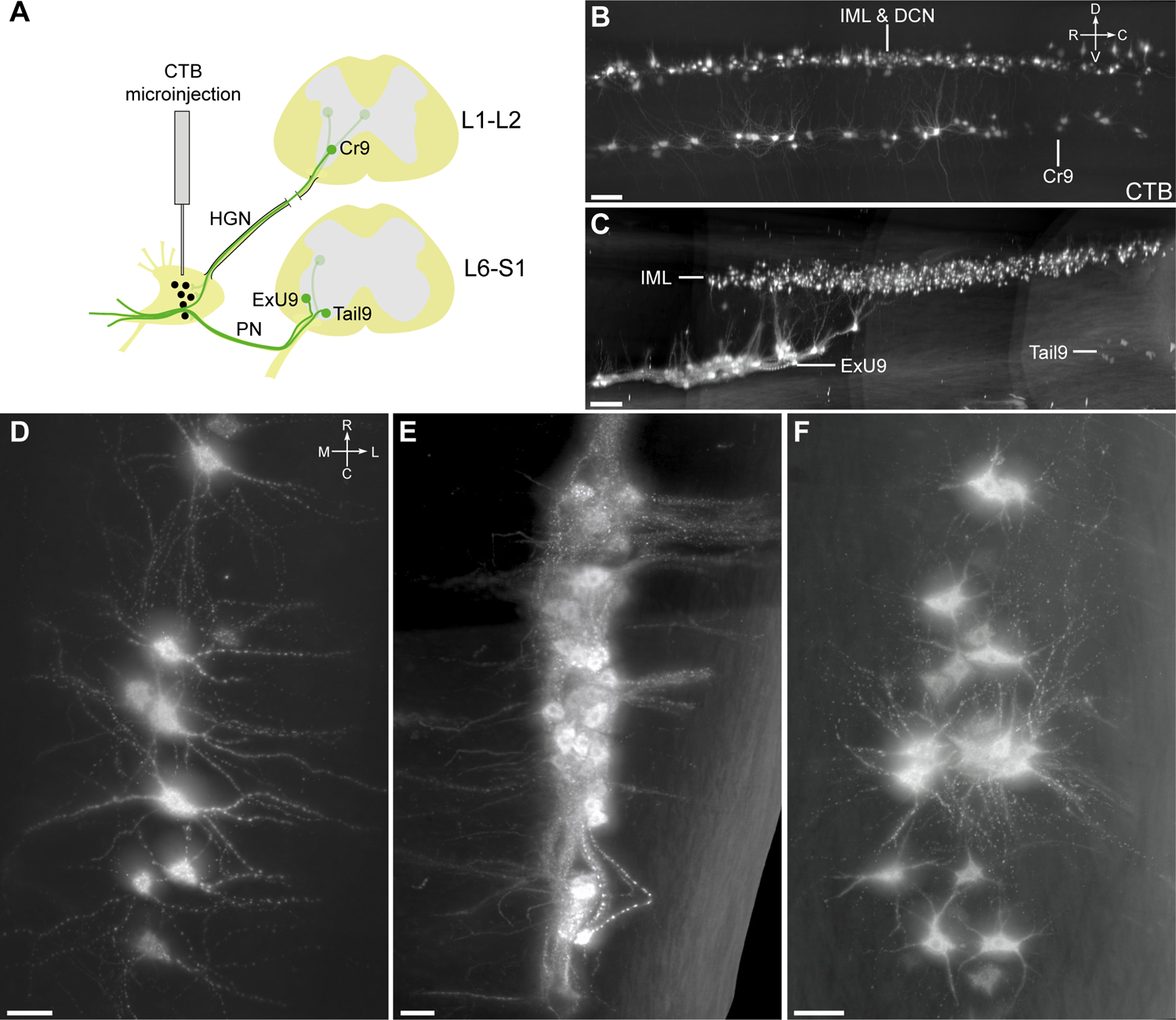
Somatomotor projections via the major pelvic ganglion and visceral nerves, revealed by cholera toxin subunit B retrograde tracing (CTB). ***A***, Schematic representation of CTB microinjection into the rat major pelvic ganglion with resulting retrograde labelling in motor neuron populations in lumbosacral spinal cord. The inferred route of this trajectory via the hypogastric and pelvic nerves is indicated. Sagittal view of CTB-positive motor neurons in ***B***, L1-L2 and ***C***, L6-S1 cleared spinal cord (male). Horizontal virtual slices of CTB-positive motor neurons in ***D***, cremaster (Cr9), ***E***, urethral rhabdosphincter (ExU9), and ***F***, levator ani and tail (Tail9) motor nuclei. B-E, male; F, female. HGN, hypogastric nerve; PN, pelvic nerve; IML, intermediolateral nucleus; DCN, dorsal commissural nucleus. Scale bars: B&C 200 μm, D-F 50 μm.

In performing this analysis, we identified an error in the spinal cord atlas of Watson et al. (2009, 2021) as the nucleus containing the urethral sphincter motor pool (ExU9) is shown in the dorsomedial MN group close to the midline (Fig. 5) whereas multiple studies locate the ExU9 motor pool in the dorsolateral MN group on the lateral margin of the ventral horn (Schrøder, 1980; Vizzard et al., 1995; Nadelhaft and Vera, 1996, 2001). Based on this evidence we have used a revised corrected nomenclature in this report, including all figure annotations, and not the original annotations shown in the atlas schematic (Table 3).

**Table 3:** Nomenclature and muscle targets of lumbosacral somatic motor neurons.

| **Romanes<sup>1</sup>**<br>**nomenclature** | **Watson<sup>2</sup>**<br>**nomenclature** | **Innervation targets** |
| --- | --- | --- |
| Ventral | Ax9<br>Tail9 | Axial muscle,<br>Tail, levator ani |
| Lateral | Hm9<br>Gl9<br>CFI9<br>CEx9 | Hamstring muscles<br>Gluteal muscles<br>Crural flexor<br>Crural extensors |
| Dorsolateral | ExU9<br>Hm9<br>Gl9 | Urethral rhabdosphincter<br>Hamstring muscles<br>Gluteal muscles, bulbocavernosus |
| Retrodorsolateral | Pes9 | Distal crural muscles |
| Dorsomedial | ExA9 | Anal rhabdosphincter |

<sup>1</sup>Romanes (1953), Schrøder (1980)<sup>2</sup>Watson et al. (2009, 2021)

### Visceral and somatic MN pools in the lumbosacral transition area

The motor system in the lumbosacral transition area within spinal cord segments L5-S2 has unique neuroanatomical organizational features that support control of sexually dimorphic visceral-somatic motor function. Here, pelvic VMNs (parasympathetic preganglionic) are collocated with the SMN pools that target striated muscles relevant to urogenital and lower bowel function previously identified by neural tracing (Schrøder, 1980). We exploited the capacity of our 3D imaging of the intact spinal cord and analysis pipeline to visualize specific groups of dorsomedial, ventral, dorsolateral and retrodorsolateral MNs and to show how these MN pools in lumbosacral spinal cord organize in 3D space (Fig. 4). Images of the L5 to S2 segments were annotated to identify the preganglionic VMNs of the IML of the parasympathetic system and the five SMN groups previously mapped by neural tracing (Schrøder, 1980). These can be rotated and viewed from any perspective in the original 3D datasets provided but are shown here as orthogonal projections from conventional transverse (Fig. 4A), horizontal (Fig. 4B) and sagittal (Fig. 4C) viewpoints. A schematic is also provided to summarize the relative rostrocaudal positioning and overlap of MN pools in the lumbosacral transition area (Fig. 4D).

**Figure 4.**
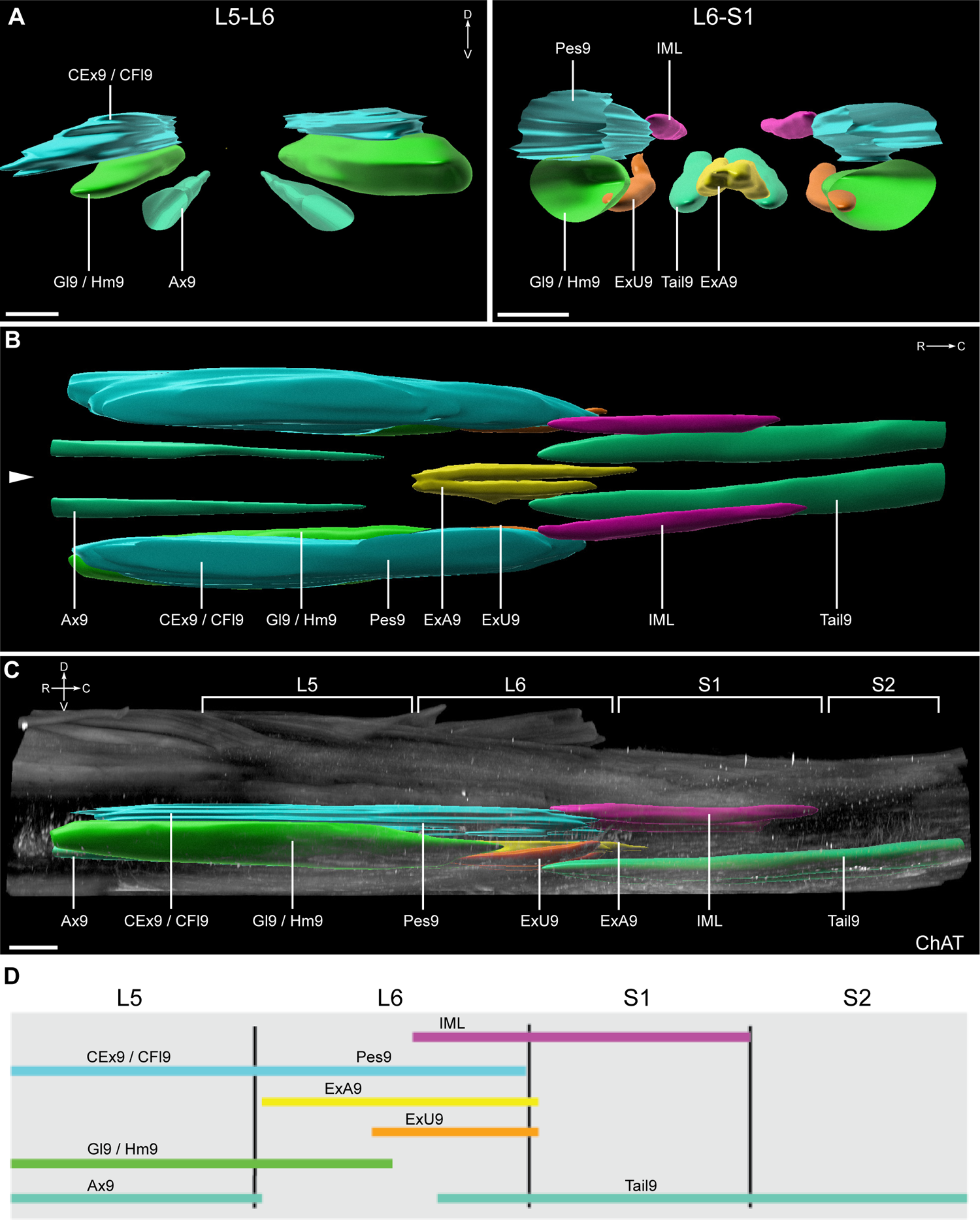
Three-dimensional mapping of lumbosacral motor pools. ***A***, Transverse view of isolated segmented motor pools in rostral (L5-L6) and caudal (L6-S1) lumbosacral spinal cord. ***B***, Horizontal visualization of the segmented motor pools in lumbosacral spinal cord. Midline indicated by arrowhead. ***C***, Orthogonal projection in sagittal view of lumbosacral spinal cord (male) with segmented motor pools superimposed on immunolabelling for choline acetyltransferase (ChAT). ***D***, Schematic of the rostrocaudal distribution of motor and premotor nuclei from L5 to S2. Mapping based on datasets from ribbon scanning confocal microscopy. CFl9, crural flexors; CEx9, crural extensors; Gl9, gluteal; Hm9, hamstring; Ax9, axial; Pes9, foot; ExU9, urethral rhabdosphincter; ExA9, anal rhabdosphincter; IML, intermediolateral nucleus; Tail9, tail. Scale bar: 500 μm (applies to all panels).

Further appraisal of this 3D map showed that L6 was an especially complex segment of the spinal cord, where the most rostral half contained no VMNs but in the caudal half the VMNs were located near four quite distinct groups of SMNs (ExU9, ExA9, Pes9, Tail9). In contrast, the VMNs of S1 extended along the complete segment and were accompanied by only one group of SMNs (Tail9) (Fig. 4D). This anatomy creates a major challenge for selecting a single representative transverse section of L6 to demonstrate its structure in a conventional 2D atlas (Watson et al., 2009, 2021). Moreover, this current spinal cord atlas is based on Wistar rats, whereas many studies are performed on Sprague-Dawley rats where strain differences have been reported in the rostrocaudal location of the parasympathetic preganglionic neurons (Pascual et al., 1989); L6-S1 in Sprague-Dawley, S1-S2 Wistar).

To account for strain differences and compare our 3D data to the 2D atlas maps from Wistar rats, we produced an ordered series of 24 virtual transverse slices (6 per segment) of the L5 to S2 segments in male and female spinal cord of Sprague-Dawley rats. These were immunolabelled with ChAT to identify the MN nuclei, and NeuN to label the soma of most spinal cord neurons, an approach widely used to reveal central nervous system cytoarchitecture.

Most spinal cord atlases identify MN pools as nuclei in spinal cord lamina 9 that are named after the target of the largest MN pool found at that location. The atlas of the adult rat spinal cord Watson et al. (2009, 2021) provides a map for each of the L5 to S2 segments in the lumbosacral transition area (Fig. 5, *left panel*). These maps identify the IML (or SPSy) and eight lamina 9 nuclei in locations corresponding to the five MN groups identified by neural tracing (Table 3).

Using the series of virtual slices from our 3D male rat datasets, we identified the four virtual slices that most closely aligned with the atlas maps of the male spinal cord and show their actual locations on a horizontal view of the lumbosacral transition area (Fig. 5, *right panel*). This revealed mismatches between our maps based on a 3D approach with segments defined by ventral roots (Sprague-Dawley rats) and the segmental representative 2D sections selected for the atlas (Wistar rat). First, our virtual slice resembling the 2D atlas view of L6 was located at the L5-L6 boundary. Even if there were no difference in rat strains, a section at this level would not be expected to contain preganglionic neurons. Second, our virtual slice resembling the 2D atlas view of S2 was located within S1. Together, this comparison demonstrated how the four 2D atlas maps of the lumbosacral transition area lack the spatial resolution needed to capture changes of the complex 3D topology of the MN pools that occur within the segments of the lumbosacral transition area. To provide a more granular resource for research on the lumbosacral cord, we therefore produced from our 3D datasets an ordered series of transverse virtual slices of ChAT immunolabelling in male and female rat spinal cord (Fig. 6).

**Figure 5.**
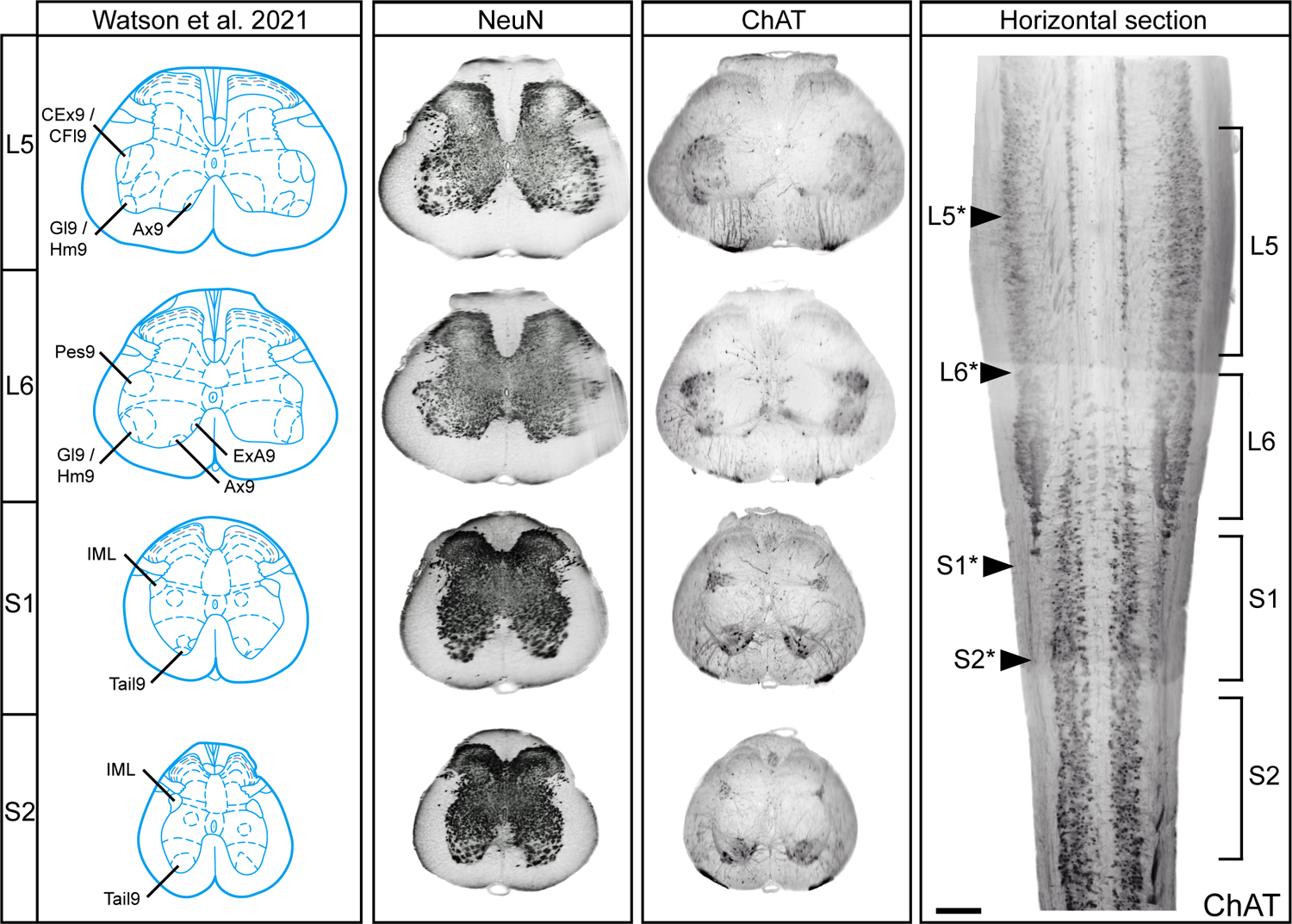
Identification of rostrocaudal position of rat spinal cord atlas sections along the lumbosacral neuraxis. Matching representative diagrams of segments L5 to S2 taken from the rat spinal cord atlas (Watson et al., 2021)to 40 μm transverse virtual slices of cleared spinal cord labelled with either NeuN or choline acetyltransferase (ChAT) (male). Rostrocaudal position of each transverse virtual slice indicated on horizontal slice (with arrows and asterisk) of same spinal cord relative to actual segment position, as determined by ventral roots. CFl9, crural flexors; CEx9, crural extensors; Gl9, gluteal; Hm9, hamstring; Ax9, axial; Pes9, foot; ExU9, urethral rhabdosphincter; ExA9, anal rhabdosphincter; IML, intermediolateral nucleus; Tail9, tail. Scale bar: horizontal section, 500 μm.

**Figure 6.**
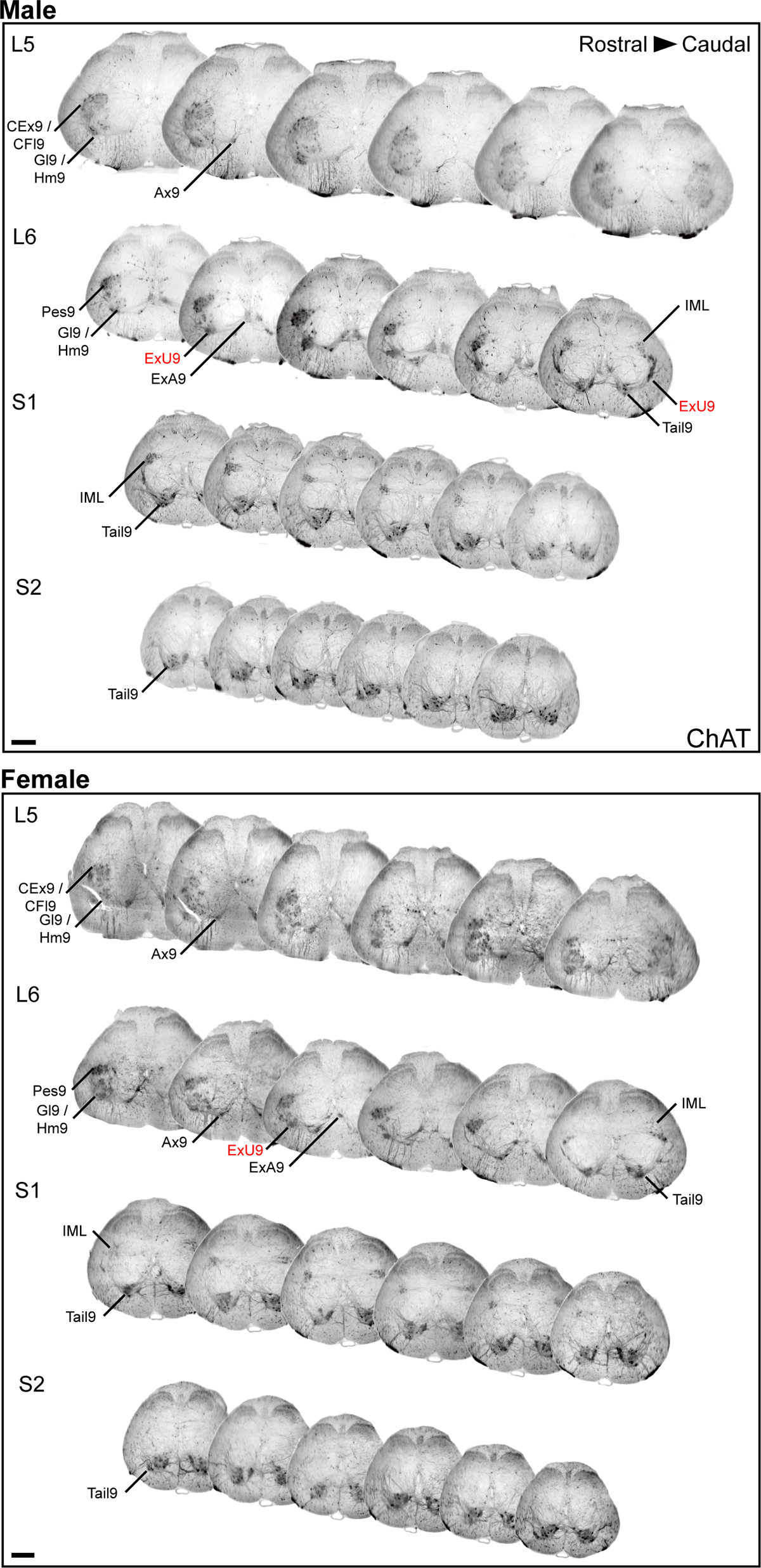
Comprehensive visualization of motor neuron topography in male and female rat lumbosacral spinal cord. Transverse virtual slices extracted at regular intervals across the 3D lumbosacral spinal cord dataset; six slices shown per segment. Spinal cord immunolabelled with choline acetyltransferase (ChAT). Labels in red indicate a correction compared to the Watson et al. (2021) rat spinal cord atlas based on the literature and our observations. CFl9, crural flexors; CEx9, crural extensors; Gl9, gluteal; Hm9, hamstring; Ax9, axial; Pes9, foot; ExU9, urethral rhabdosphincter; ExA9, anal rhabdosphincter; IML, intermediolateral nucleus; Tail9, tail. Scale bar: 500 µm (applies to both panels).

### Dendritic fields of parasympathetic VMNs reveal sexual dimorphism

Preganglionic VMNs have a characteristic multipolar morphology with long dendrites, which in a single neuron can span the distance from the midline to the lateral edge of the white matter tracts (Nadelhaft and Booth, 1984; Anderson et al., 2009; Peddie and Keast, 2011). The relative density of somatic versus dendritic inputs has not been quantified but large multipolar CNS neurons typically receive significantly more dendritic than somatic input (Nadelhaft and Booth, 1984; Morgan et al., 1993; Morgan and Ohara, 2001) This is consistent with projections of supraspinal inputs from the pontine micturition center to the soma of lumbosacral (parasympathetic) preganglionic MNs and following the medial dendrites out to the midline (Valentino et al., 1995, 1999; Keller et al., 2018). This illustrates the important functional role of the dendritic field volume or “peri-IML” in receiving and integrating spinal and supraspinal input.

As CTB labelling identifies a much greater proportion of the MN dendritic tree than ChAT immunohistochemistry, we explored data from our MPG neural tracing experiments to assess how the dendrites of parasympathetic preganglionic neurons project in 3D space (Figs. *7-9*, Media 4 [https://doi.org/10.26188/25484332.v1]). Transverse views of the L6-S1 IML (Fig. 7) showed how the dendrites have three preferred orientations: *medial*, often extending into the sacral dorsal commissural nucleus (SDCom) and close to the midline; *ventrolateral* extending deep into the ventral horn; and *lateral* into the surrounding spinal white matter tracts. These orientations are demonstrated in horizontal and sagittal views in Fig. 8. Previous studies tracing single IML neurons in rodents and other species also report long rostrocaudal dendrites (Armstrong et al., 1983; Barber et al., 1984) These were difficult to view within the IML where they run between the strongly fluorescent somata but were shown to extend rostrally for some distance past the end of the preganglionic neuron column (Fig. 8A).

**Figure 7.**
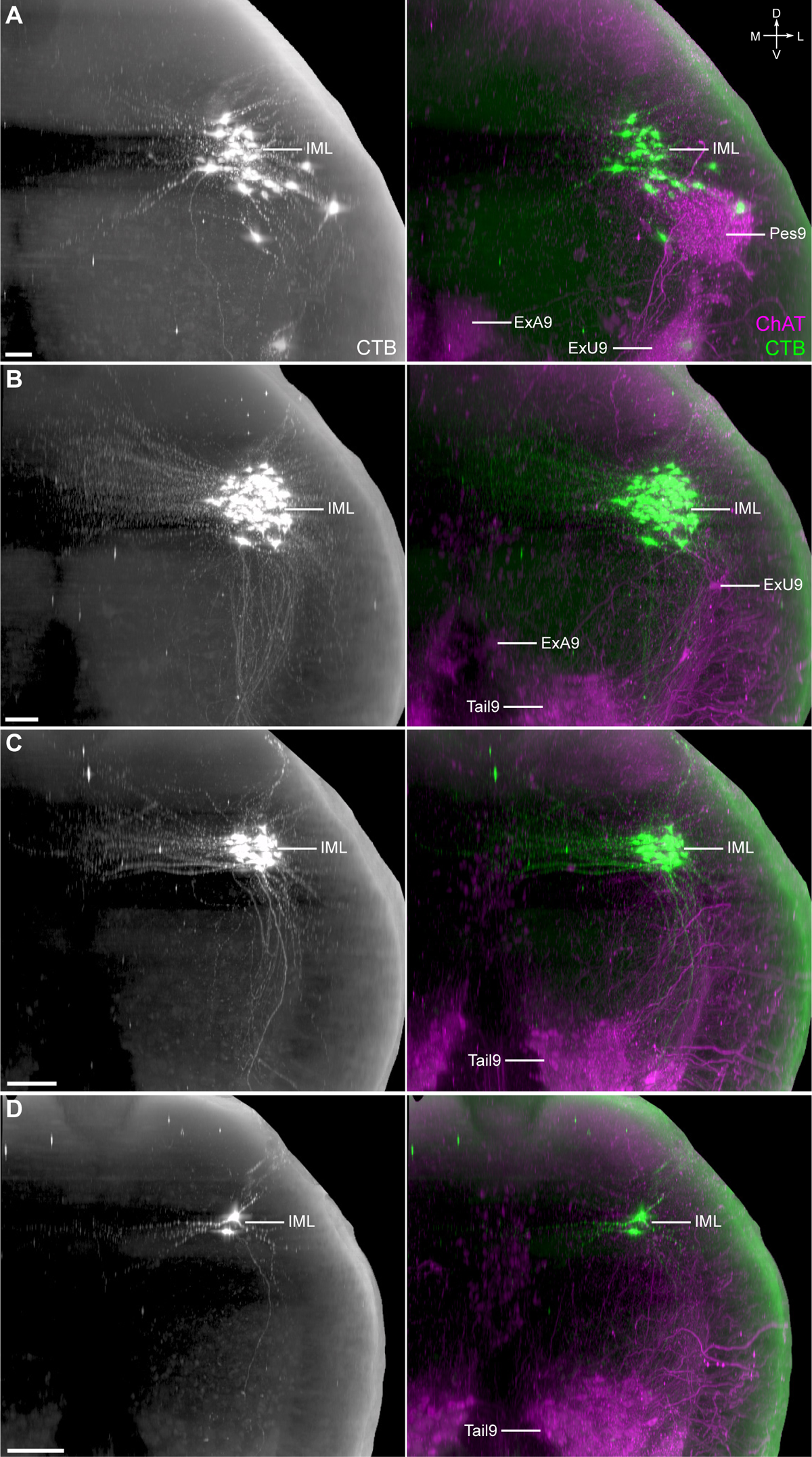
Dendritic field visualization achieved through retrograde filling of lumbosacral preganglionic neurons. Transverse virtual slices (100 μm) of cleared spinal cord from ***A***, caudal L6, ***B***, rostral S1, ***C***, middle S1, and ***D***, caudal S1. Spinal cord immunolabelled for cholera toxin subunit B (CTB) and choline acetyltransferase (ChAT). IML, intermediolateral nucleus; Pes9, foot; ExU9, urethral rhabdosphincter; ExA9, anal rhabdosphincter; Tail9, tail. Scale bar: 100 μm.

**Figure 8.**
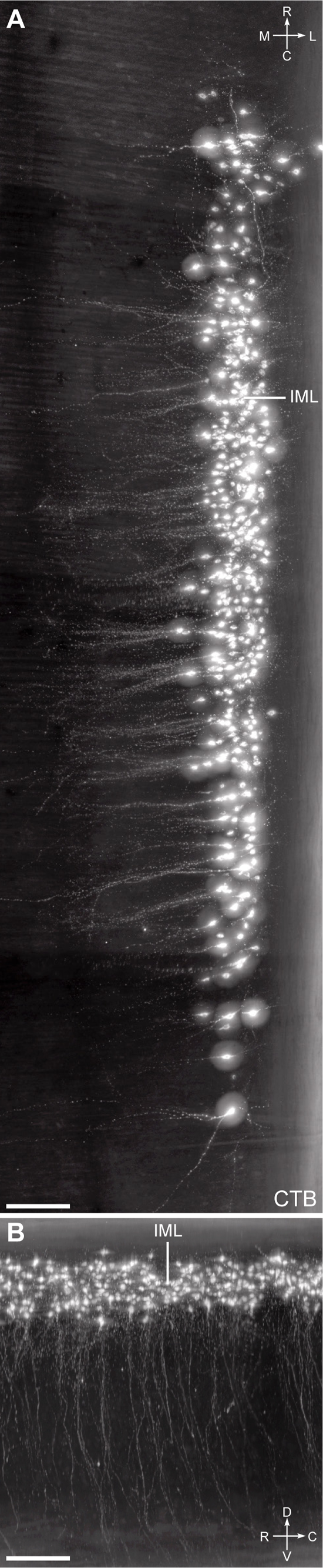
Horizontal and sagittal views of dendritic field originating from sacral preganglionic neurons. 200 μm thick virtual slices in ***A***, horizontal and ***B***, sagittal orientations, focussing on the intermediolateral nucleus (IML). Spinal cord immunolabelled for cholera toxin subunit B (CTB). Intensity of signal has been digitally increased to demonstrate dendrites in CTB+ neurons. Scale bar: 200 μm.

**Figure 9.**
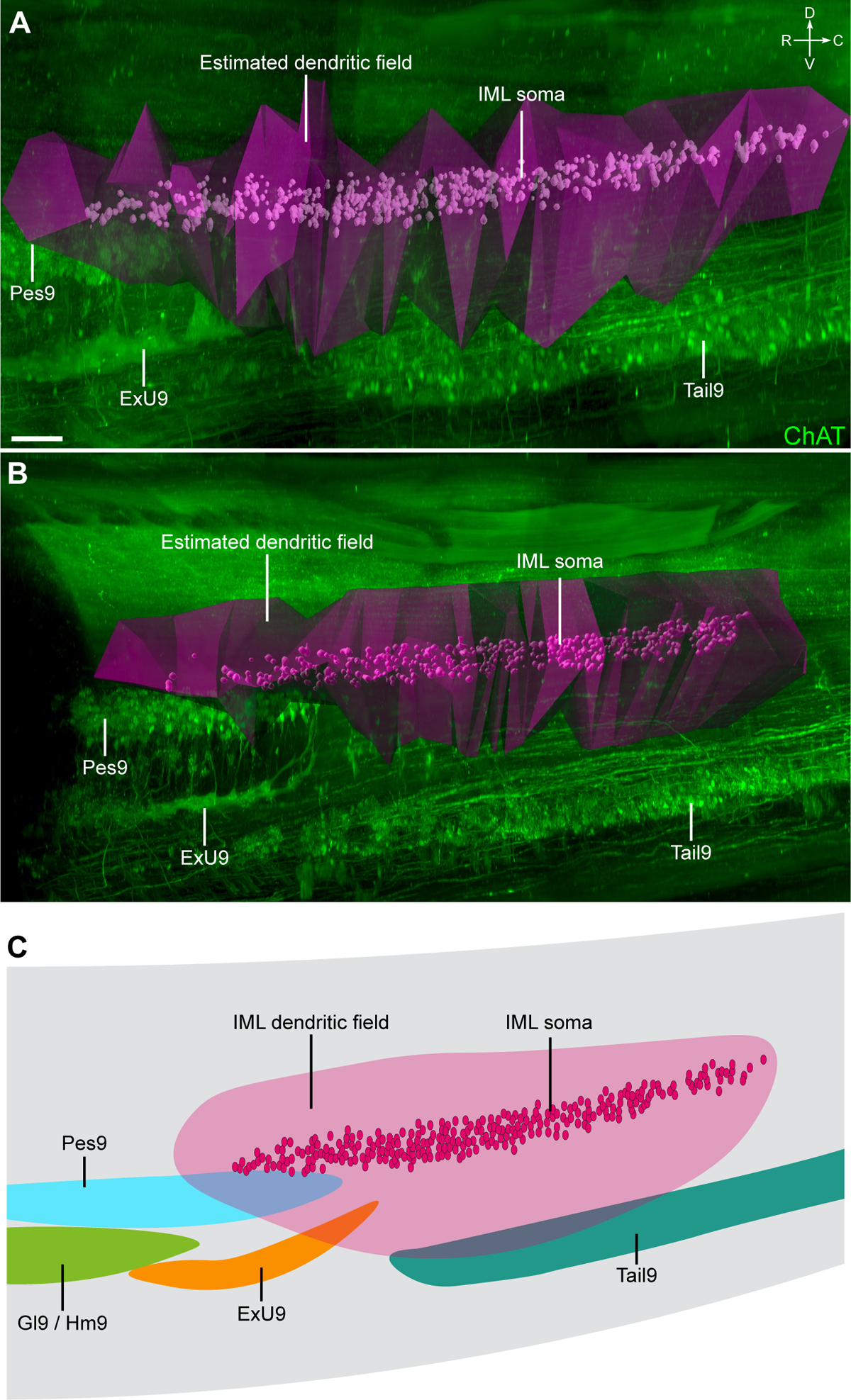
Estimated dendritic field representation of lumbosacral preganglionic neurons in the 3D context of the cleared lumbosacral spinal cord. 3D volumes were generated according to the distribution of cholera toxin subunit B in the dendrites of preganglionic neurons located in the intermediolateral column of L6-S1 spinal cord in ***A***, male, and ***B***, female rats. Spinal cord immunolabelled for choline acetyltransferase (ChAT). ***C***, Schematic representation of dendritic field of lumbosacral preganglionic neurons in relation to nearby motor pools. Pes9, foot; ExU9, urethral rhabdosphincter; ExA9, anal rhabdosphincter; Tail9, tail. Scale bars: 100 μm (applies to both panels).

The strong labeling of CTB^+^ lumbosacral (parasympathetic) preganglionic neurons encouraged us to visualize and quantify their dendritic field volume. The CTB immunofluorescence in dendrites was punctate, similar to that recently reported in central axons of sacral spinal sensory neurons. We used the *Spots* function in Imaris to segment the fluorescent CTB^+^ puncta to obtain XYZ coordinates of a cloud of points outlining preganglionic neurons and their dendrites, and then analyzed with a custom code that used a 3D composite convex hull to fit a surface around the point cloud (https://gitlab.unimelb.edu.au/lab-keast-osborne-release/3d-points-volume-estimation). A comparison of the total dendritic field volume of CTB^+^ lumbosacral preganglionic neurons in three male (Fig. 9A) and female (Fig. 9B) spinal cords identified a sex difference, being significantly larger in males (1.201 ± 0.188 mm^3^ vs 0.624 ± 0.099 mm^3^; difference = 0.578 ± 0.219 mm^3^, 2-tailed *t-*test: *P =* 0.0388).

Many MNs in the spinal cord contribute to dendritic bundles. These are large groups of closely apposed parallel dendrites that can be electronically coupled by dendro-dendritic gap junctions. It is hypothesized that dendritic bundles have a role in synchronizing activity within or across functional pools of MNs (Matthews et al., 1971; Personius et al., 2007; Bautista and Nagy, 2014). The largest dendritic bundles are found between SMNs in the lumbosacral transition area (Roney et al., 1979; Schrøder, 1980; Bellinger and Anderson, 1987). To our knowledge there are no reports of dendritic bundles including dendrites from both SMNs and preganglionic VMNs, but our analysis of dendritic field volumes showed that dendrites from these VMNs were close to the ExU9, Pes9 and Tail9 MN pools (Fig. 9C). We further examined the spinal cords from the CTB injection study, where both lumbosacral preganglionic neurons and specific pools of somatic motor neurons were labelled from the MPG. Higher magnification images (Fig. 10, Media 5 [https://doi.org/10.26188/25484764.v1]) showed there were many CTB^+^ dendrites in bundles linking the lumbosacral IML and ExU9.

**Figure 10.**
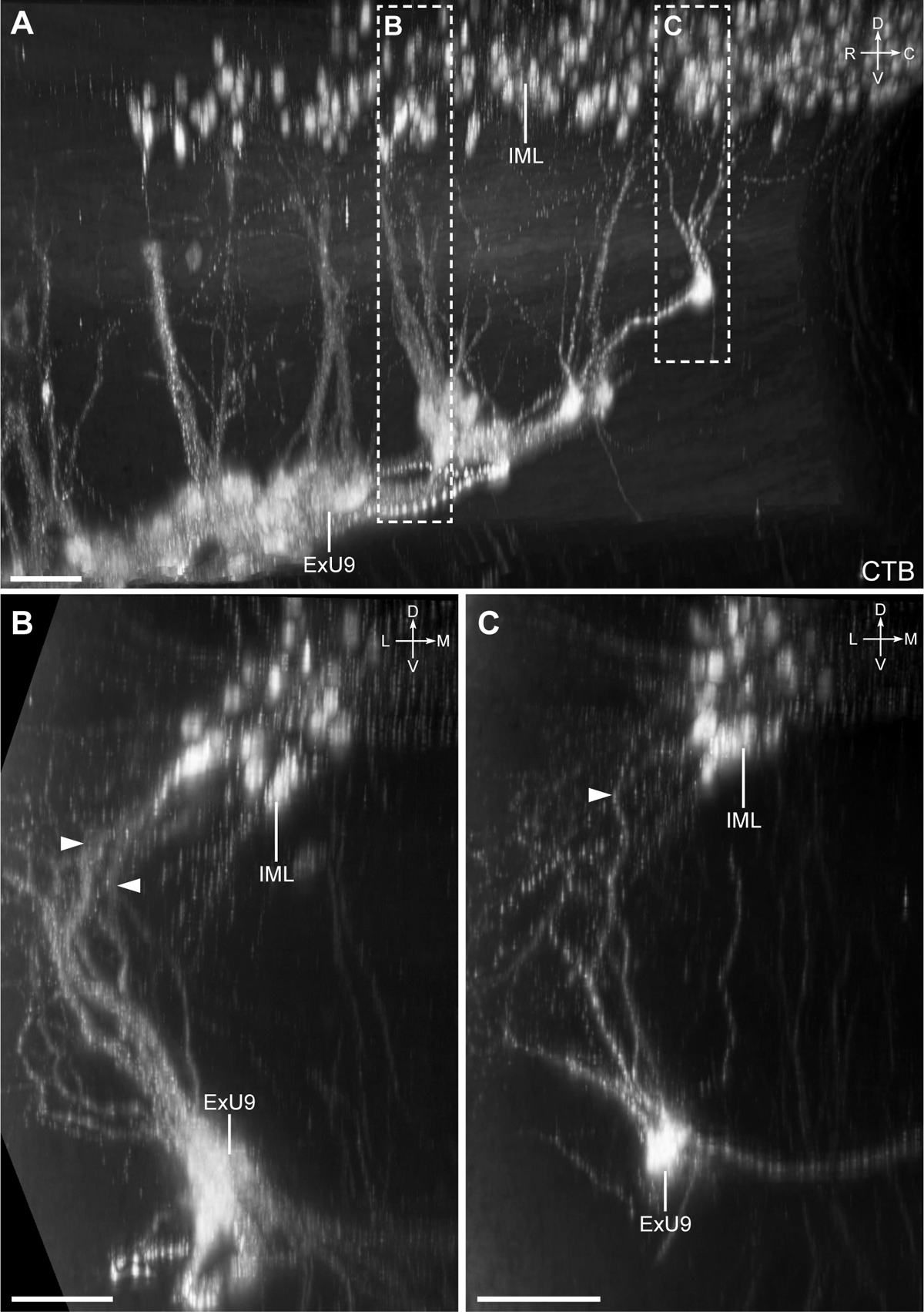
Close association between dendrites of lumbosacral visceral (preganglionic) and somatic motor neurons (MNs) identified with cholera toxin subunit B (CTB) retrograde filling. CTB was microinjected into the major pelvic ganglion. ***A***, Sagittal view of urethral rhabdosphincter (ExU9) MN dendrites projecting dorsally towards visceral MNs of the intermediolateral nucleus (IML) in cleared spinal cord (L6). ***B*** and ***C***, ExU9 dendrites (indicated by arrows) in close proximity to less brightly labelled dendrites of the IML neurons, visualized in 100 μm transverse virtual slices. Scale bars: 100 μm (applies to all panels).

### Sexual dimorphism of rhabdosphincter SMNs

Having identified a sex difference in the dendritic field volume of parasympathetic preganglionic neurons and finding evidence of their dendritic bundles with ExU9, we next investigated if there was sexual dimorphism in the urethral rhabdosphincter MN pool (ExU9). This has been previously examined in reconstructions from histological sections (Breedlove and Arnold, 1981; Jordan et al., 1982; Sengelaub et al., 1989; Tobin and Payne, 1991). In our 3D datasets, ChAT immunolabelling showed ExU9 was larger and extended further caudally in males than females (Fig. 11A). However, the intensity of the ChAT fluorescence in the dendritic bundles that extend from ExU9 made it difficult to count the less intensely labelled somata of the MNs. To address this, we reanalyzed an open dataset (Fuller-Jackson et al., 2021a) containing images of an ordered series of sections taken through the lumbosacral spinal cord in adult female and male Sprague-Dawley rats. This visualization of ChAT^+^ neurons in sections allowed us to segment individual MN somata from the raw signal. Plotting relative nucleus area along the rostrocaudal axis of the nucleus confirmed ExU9 (Fig. 11B, C) was larger and extended over a longer distance in males (ExU9 transverse area: male = 36,627 ± 13,885 µm^2^, n =6 versus female = 21,128 ± 3895 µm^2^, n = 5; two-tailed *t-*test, *P* = 0.026). Similar analyses determined ExA9 (Fig. 11D, E), the nucleus that targets the anal rhabdosphincter, also was larger in males (males, 16,242 ± 6040 µm^2^, n = 6 versus females, 6191 ± 2486 µm^2^, n = 6; two-tailed *t-*test, *P* = 0.003).

**Figure 11.**
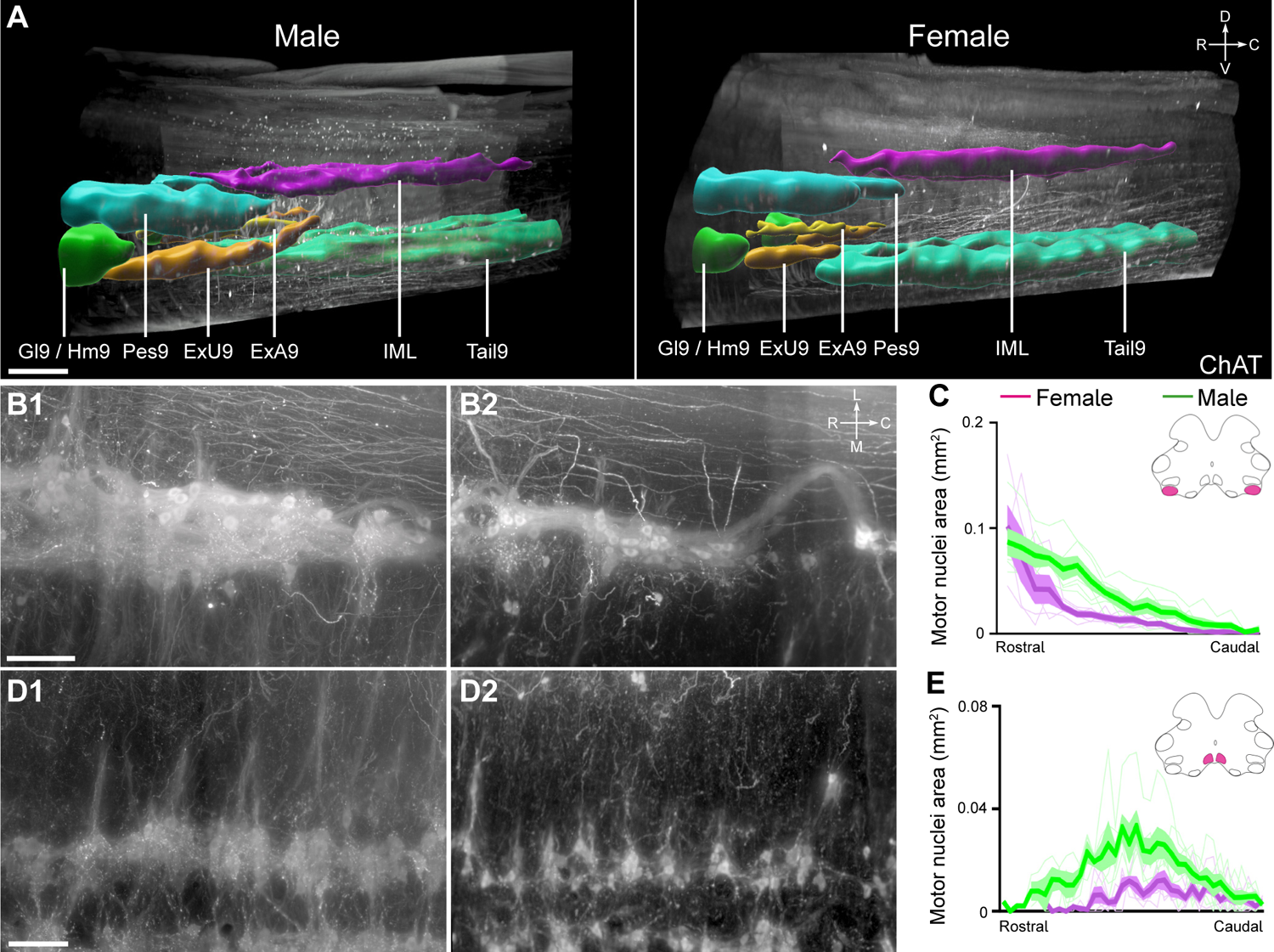
Sexual dimorphism of motor nuclei L6-S1. ***A***, Sagittal view of cleared lumbosacral spinal cord (L6-S1) from male and female rat, immunolabelled with choline acetyltransferase (ChAT) and motor nuclei segmented. ***B***, Horizontal virtual slice of urethral rhabdosphincter motor neurons (ExU9) of male (B1) and female (B2) rats. ***C***, Urethral rhabdosphincter motor nucleus area from rostral to caudal L6 measured in transverse cryosections of male (n = 6) and female (n = 5) rats. ***D***, Horizontal virtual slice of anal rhabdosphincter motor neurons (ExA9) from male (D1) and female (D2) rats. ***E***, Anal rhabdosphincter motor nuclei area from rostral to caudal L6 measured in transverse cryosections of male (n = 6) and female (n = 6) rats. C and E, lines are mean, standard error of the mean, and individual traces contouring the relevant MN area on each section. Gl9, gluteal; Hm9, hamstring; Ax9, axial; Pes9, foot; IML, intermediolateral nucleus; Tail9, tail. Scale bars: 500 μm (A), 100 μm (B, D).

### Distinctive cellular and subcellular features of MN pools in the lumbosacral visceral-somatic transition zone

Light sheet microscopy provided a unique opportunity to build a true 3D map of the lumbosacral MN pools and establish their mesoscale and macroscale relationships. These large 3D datasets were also of sufficient resolution to identify and compare specific microscale (cellular and subcellular) features of each MN pool, including aspects of their somata, proximal dendrites and axon projections. Here we have aggregated our descriptive findings for each of the MN pools, integrating observations illustrated in several previous Figures and extended in Figs. 12 and 13 and Movies (Media 4 [https://doi.org/10.26188/25484332.v1], Media 5 [https://doi.org/10.26188/25484764.v1], Media 6 [https://doi.org/10.26188/25484788.v1]. Together these revealed differences across groups, in agreement with previous observations from histological sections but also identifying novel features. These have been grouped according to the MN pool.

**Figure 12.**
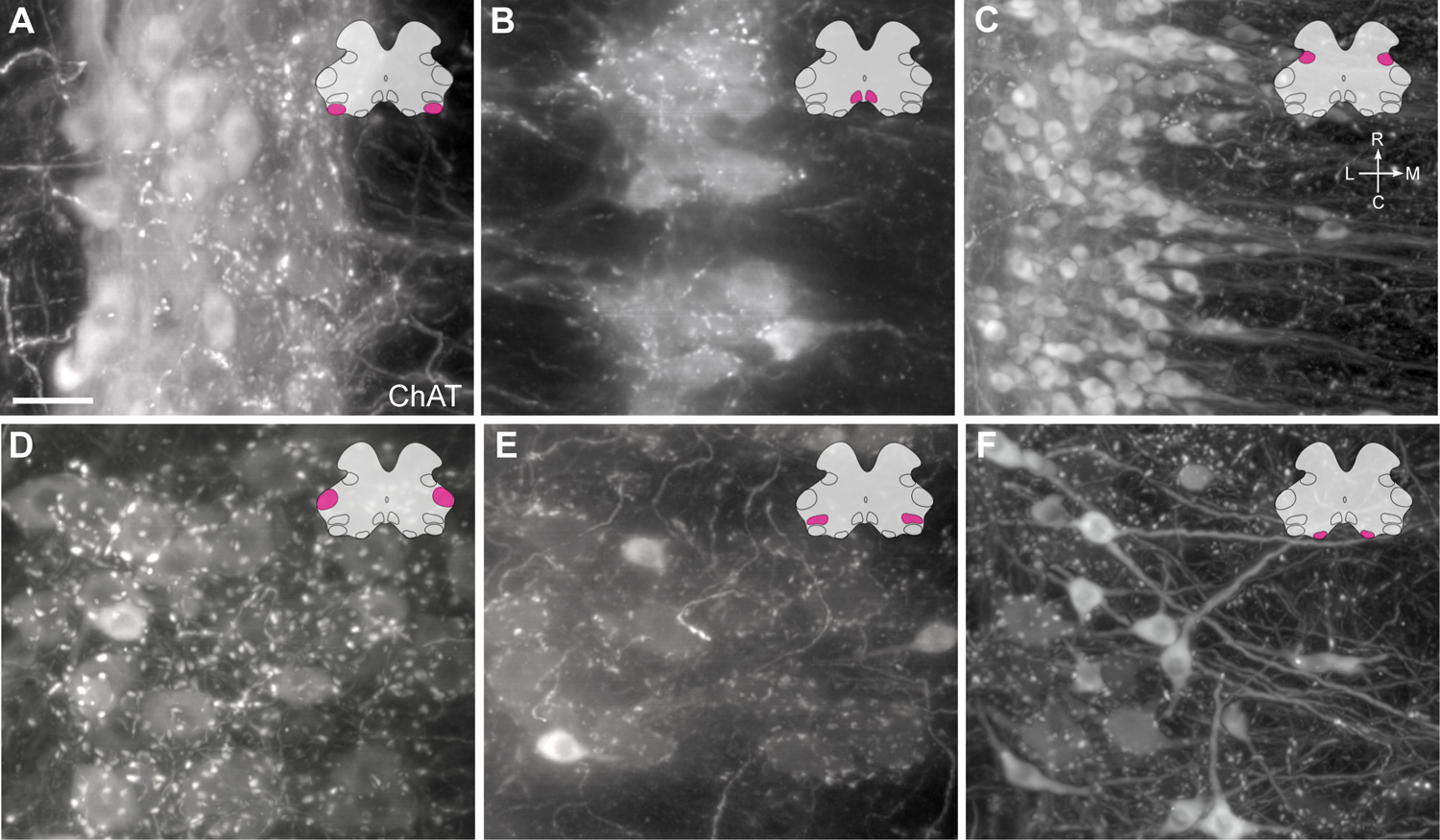
Motor neuron morphology in the lumbosacral spinal cord. Horizontal virtual slices of ***A*,** urethral rhabdosphincter (ExU9), ***B,*** anal rhabdosphincter (ExA9), ***C,*** parasympathetic preganglionic (IML), ***D,*** foot (Pes9), ***E***, hamstring and gluteal (Hm9 / Gl9), and ***F,*** levator ani and tail (Tail9) motor nuclei in the cleared lumbosacral spinal cord (L6-S1, male rat). Spinal cord immunolabelled for choline acetyltransferase (ChAT). Scale bar: 50 μm.

**Figure 13.**
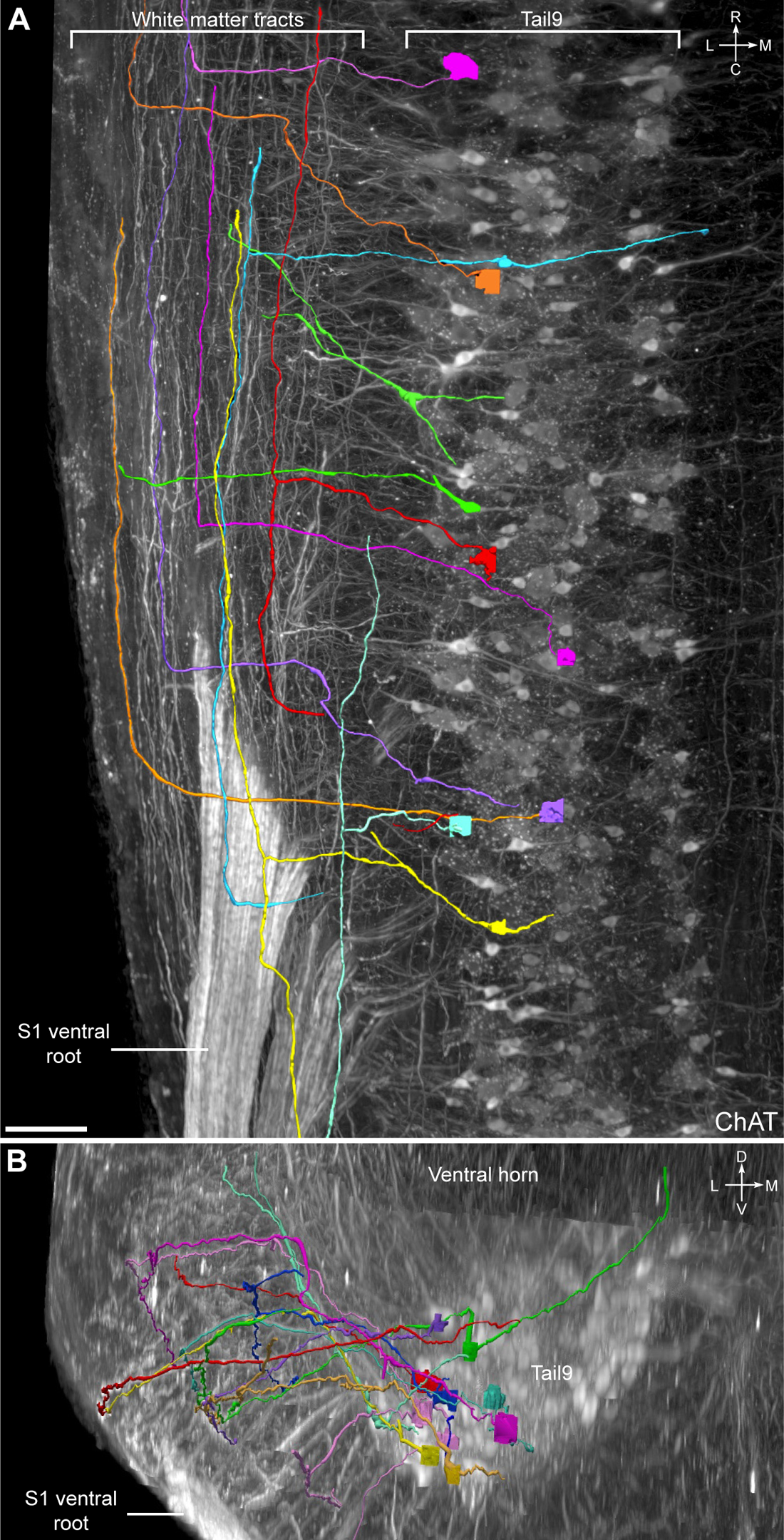
Sacral somatic motor neurons (SMNs) possess intersegmental axonal projections. ***A***, Horizontal view of the levator ani and tail SMNs (Tail9) and proximal white matter tracts in cleared S1 spinal cord with individual MN axons traced and indicated by different colours. ***B***, Zoomed-in horizontal view highlighting a SMN axon projecting into the white matter tracts. ***C***, Transverse view of ventral horn of the same cleared S1 spinal cord. ChAT, choline acetyltransferase. Scale bars: 100 μm.

### Parasympathetic (preganglionic) VMNs

The most rostral neurons in the parasympathetic IML appeared dorsomedial to the Pes9 in mid-L6 (Figs. 4, 9). Across the caudal sections of L6, the IML increased in size and neuron number, taking up the entirety of the intermediolateral region in lamina V/VI. This column of VMNs continued through S1 but did not extend into the S2 segment. At higher magnification, the VMNs formed a dense column of predominantly spindle-shaped multipolar neurons, which had smaller somata and were less rounded than SMNs (Fig. 12C). Dendrites were only partly revealed by ChAT-immunofluorescence but could be seen extending medially towards the central canal and running longitudinally between somata in the IML column; these were revealed more clearly by the retrograde tracer, CTB (Figs. 7-9, Media 4 [https://doi.org/10.26188/25484332.v1]). Proximal dendrites extending laterally into the white matter tracts were also clearly detectable with CTB but not ChAT. No large ChAT^+^ C-boutons, characteristic of typical alpha MNs, were associated with the IML.

### Urethral rhabdosphincter (ExU9)

ExU9 comprises the motor nucleus innervating the urethral rhabdosphincter and ischiocavernosus muscle (Table 3), which first appears adjacent to the ventrolateral edge of Gl9/Hm9; this corresponds to the second digital slice of L6 (Fig. 6). This is sometimes referred to as Onuf’s nucleus. By the caudal end of L6, ExU9 had reduced in size to only one to three neurons per nucleus per slice. Proceeding rostrocaudally through ExU9, the MNs gradually shifted dorsally to occupy the most lateral edge of the ventral horn and progressively showed a higher density of dorsally orientated dendritic bundles.

Although the ExU9 somata did not extend to S1, their dendritic bundles were visible in the rostral slice of S1. Dendritic bundles of ExU9 neurons extended rostrocaudally, as well as laterally and dorsally along the ventral boundary of the grey matter, with no major bundles extending dorsomedially into the deeper grey matter regions such as towards lamina X (Media 5 [https://doi.org/10.26188/25484764.v1]). Higher magnification views showed the heterogeneity across ExU9 in the distribution of c-boutons associated with somata and the density of dendritic bundles surrounding different regions of ExU9, with some ExU9 neurons have a dense supply and others having none (Fig. 12A).

### Anal rhabdosphincter (ExA9)

MNs of ExA9 supply the anal rhabdosphincter and bulbocavernosus muscles (Table 3). ExA9 is also referred to as the spinal nucleus of the bulbocavernosus (Sengelaub and Forger, 2008). In rostral L6, ExA9 appeared in the most medial portion of the ventral horn, immediately ventral to the central canal and lamina X (Fig. 6). ExA9 increased in size from rostral L6 until the mid-point (digital slices three and four of L6), after which the nuclei decreased in size until rostral S1 wherein only the ChAT^+^ dendrites of the motoneurons were visible (Fig. 6). Inspection at higher magnification demonstrated the dense meshwork of dendritic bundles surround the ExA9 MNs (Fig. 12B), although this was overall less dense than in ExU9. These bundles projected laterally in strikingly regular intervals across the rostrocaudal neuraxis (Fig. 11D).

### Lower limb (Pes9, Gl9 and Hm9)

In rostral L6, three MN nuclei innervate muscles of the lower limb; Pes9 projects to the distal crural muscle, Gl9 to the gluteal muscles, and Hm9 to the hamstring muscles (Table 3). Located in the dorsolateral corners of the ventral horn, Pes9 was a compact aggregate of predominately large, stellate and medium-sized round MNs (Figs. 6, 12D). Small, intensely labelled spindle-shaped neurons were also visible (Fig. 12D). There were no visible bundles of dendrites, however c-boutons were present around many large and medium-sized MNs (Fig. 12D). By caudal L6, Pes9 was much smaller, with few MNs remaining and none appearing in S1 (Fig. 6). In the ventrolateral corner of the ventral horn in rostral L6, Gl9 and Hm9 MNs also comprised large, medium and small MNs of stellate, round and spinal morphology, respectively (Fig. 12E). Compared to MNs of Pes9, Gl9 and Hm9 MNs were less densely packed (Figs. 6, 12D). Halfway through L6 (the fourth slice), Gl9 and Hm9 ended, to be replaced completely by ExU9 MNs (Fig. 6).

### Axial (Ax9)

Ax9 projects to the axial muscles and is located in the most ventromedial aspect of the L5 ventral horn (Fig. 6). This is a small nucleus, with only 1-2 neurons per slice observed in rostral L6 and ending by the third slice. At this point, it was replaced by Tail9.

### Levator ani and tail (Tail 9)

The levator ani and tail muscles are innervated by the Tail9 nucleus, which appeared in the third slice of L6 in the position previously occupied by Ax9 in the more rostral slices (Fig. 6). In the fourth slice, the diameter of the Tail9 nucleus greatly increased and by the most caudal slice of L6 Tail9 was expanded in size such that throughout the S1 and S2 segments it occupied around half of the ventral horn. At higher magnification, Tail9 MNs in the ventral horn of the sacral spinal cord exhibited the greatest disparity between neuronal subtypes (Fig. 12F). Large stellate motoneurons with weak ChAT immunolabelling were surrounded by the greatest density of c-boutons. Medium-sized round motoneurons had stronger ChAT immunolabelling but fewer c-boutons around their soma. Small spindle-shaped motoneurons devoid of c-bouton contacts were the most intensely immunolabelled motoneurons.

Of the lumbosacral motor nuclei inspected at higher magnification, the axons of the Tail9 MNs were the most strongly immunolabelled. Many could be followed into the white matter tracts, turning 90° to project rostrally amidst a column of ChAT^+^ axons in the ventrolateral white matter tract (Fig. 13, Media 6 [https://doi.org/10.26188/25484788.v1]). Tracing these axons in Neurolucida 360 showed that all axons travelling via the white matter reached the L6-S1 boundary, presumably continuing to rostral segments. Motoneurons that possessed these axonal projections into the white matter were of no single type, i.e., including both small spindle-shaped neurons and large neurons surrounded by c-boutons. Some axons instead turned to project caudally, however this was less frequently observed. In two cases, axons were found to bifurcate in the white matter, projecting in both directions of the neuraxis (Fig. 13A).

### A comprehensive map of lumbosacral motor innervation

The topographical organization of the SMN and VMN nuclei in the lumbosacral cord, particularly the complexity of the L6 segment, requires a more comprehensive graphical representation than is currently available. Based on our multi-scale mapping, we created a series of schematics indicating the approximate position of each of the SMN and VMN nuclei as they would appear in transverse sections (Fig. 14A). These have been colored according to the somatic muscle or pelvic organ that they regulate, in both male (Fig. 14B, C) and female (Fig. 14D) Sprague-Dawley rats.

**Figure 14.**
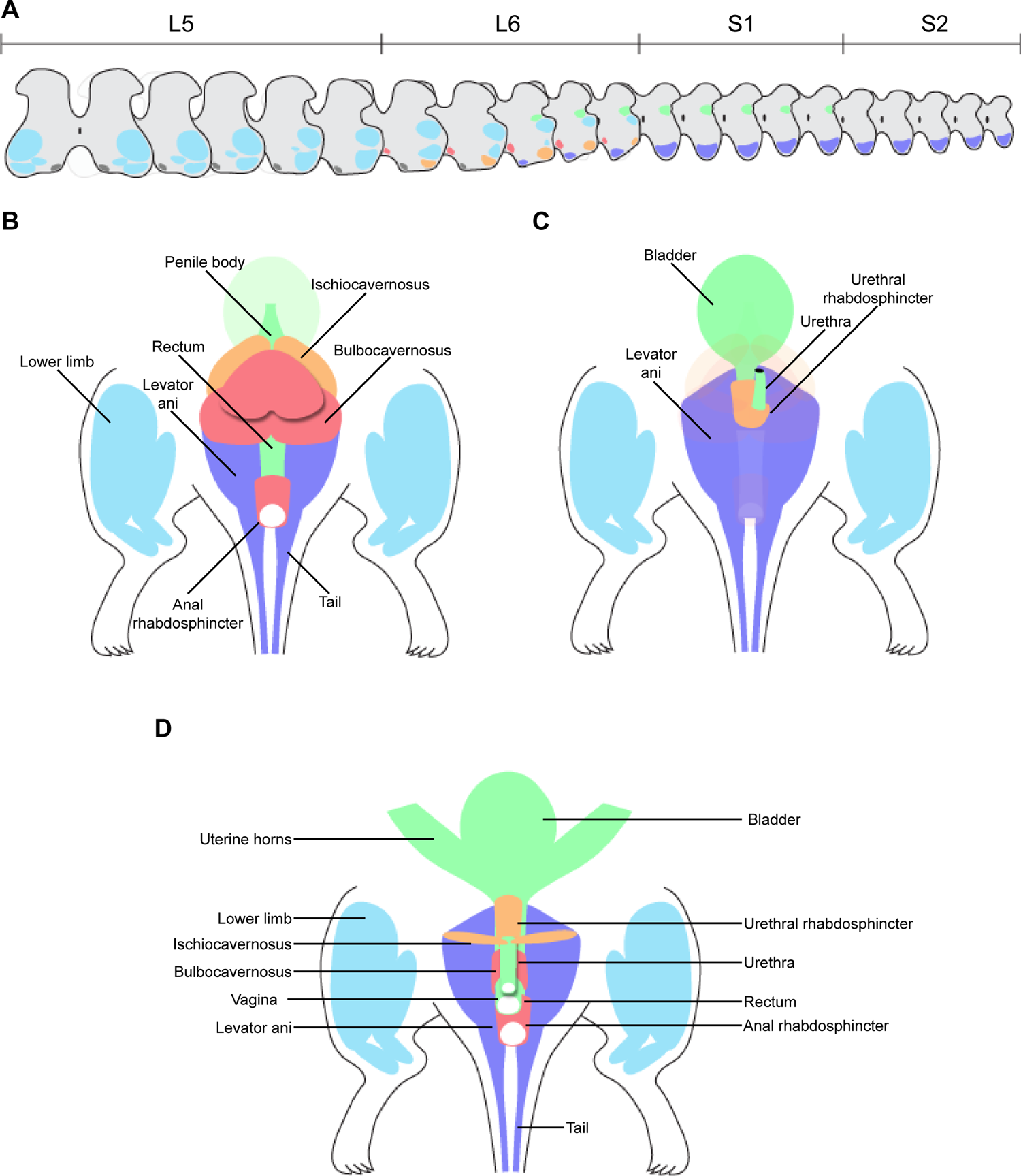
Graphical summary of lumbosacral motor nuclei topography and innervation. ***A***, Topography of motor nuclei in transverse L5 to S2 spinal cord. Colorization of motor nuclei matches that of the muscles in B-D, which illustrate the muscles and organs innervated by somatic and visceral components of the pelvic motor system. ***B*** and ***C***, Male rat from dorsal (B) and ventral (C) views. ***D***, Female rat, ventral view.

## Discussion

We applied advanced whole mount immunofluorescence 3D microscopy (Belle et al., 2017; Fuller-Jackson et al., 2021b; Blain et al., 2023) to study the pelvic MN system in the lumbosacral spinal cord of female and male rats. This highly specialized motor circuit formed by autonomic preganglionic VMNs and SMNs controls motor activity during excretory, sexual, and reproductive behaviors (Anderson et al., 2009; Holstege and Collewijn, 2009; de Groat, 2018). Unusual features include extensive morphological sex differences, motor patterns that combine visceral and somatic activity, and psychogenic activation by the brain during higher order behaviors (Breedlove and Arnold, 1981; Thor and de Groat, 2010; Hou et al., 2016; Keller et al., 2018). 3D analysis of spinal cord neuroanatomy has been limited by the technical challenges of producing reconstructions from sections, but with new technologies we could view MNs in 3D at microscopic resolution and perform the first multiscale study of macroscopic 3D organization and microscopic morphology of pelvic MNs. Our datasets will be available as a resource on an open science platform (sparc.science) to allow further exploration of the lumbosacral spinal motor system.

We first revealed the macroscopic organization of MNs with RCSM (Watson et al., 2017), using the characteristic size, morphology and location of cholinergic MNs to distinguish them from cholinergic interneurons dispersed in other spinal cord regions (Barber et al., 1984). During development, MNs segregate into longitudinal medial and lateral columns (Dasen, 2022), which postnatally are interrupted by the enlargements to produce subcolumns of morphologically distinct MNs (Barber et al., 1984). The 3D organization of the MN system in the lumbosacral spinal cord was shown by segmenting these columns and projecting digital 3D models onto raw images. This demonstrated how composition of the functionally distinct MN columns differ in upper, middle, and lower segments of the lumbosacral cord. The lateral SMN columns targeting the hindlimbs occupy the middle region at the center of the enlargement (Nicolopoulos-Stournaras and Iles, 1983). These extend into the rostral region where they overlap with the ventral SMN columns and caudal end of the sympathetic VMN columns (Anderson et al., 2009; Scott-Solomon et al., 2021; Qi et al., 2022). In the caudal region, the lateral column overlaps with a single parasympathetic VMN column and four SMN columns (Schrøder, 1980; Barber et al., 1984). The increased number and neuroanatomical complexity of these MN columns corresponds with this region being the major source of motor output to the pelvis and urogenital region (Holstege and Collewijn, 2009; de Groat, 2018).

We identified both sympathetic (L1-L2) and parasympathetic (L6-S1) pelvic VMNs by neural tracing from their primary output target, the bilateral MPG (Hancock and Peveto, 1979b, 1979a; Watkins and Keast, 1999). This confirmed previous evidence indicating major differences between the pelvic VMNs and other thoracolumbar VMNs. First, major targets of the thoracolumbar (sympathetic) IML are the paravertebral (sympathetic chain) ganglia that control cardiovascular (vasoconstrictor) function and thermoregulation (Anderson et al., 2009). We confirmed that most thoracolumbar VMNs regulating pelvic organs are instead located in a midline column, the DCN (Hancock and Peveto, 1979a, 1979b; Watkins and Keast, 1999), rather than the IML. Second, the parasympathetic VMNs located in the L6-S1 IML only target the MPG and therefore exclusively regulate visceral targets, again contrasting with a major function of the sympathetic (thoracolumbar) IML. These functional differences between thoracolumbar IML, DCN, and lumbosacral IML are relevant to recent transcriptomic classification of MN classes (Blum and Gitler, 2022). Functional specialization of the VMNs in IML above and below the lumbar enlargement correlates with their distinct transcriptome features (Alkaslasi et al., 2021; Blum et al., 2021; Liau et al., 2023). The central column of VMNs in the DCN that preferentially output to pelvic visceral motor targets have not yet undergone this type of analysis, but this is predicted to reveal neural classes different to those in the IML above and below the lumbar enlargement.

The retrograde labelling of some SMNs in ExU9, Cr9 and Tail9 by MPG microinjection of CTB indicates that specific SMN groups innervate their target muscles via an MPG trajectory, which has potential implications for surgical procedures and therapeutic neuromodulation. In cats and humans, there are multiple reports of communicating branches between the pudendal nerve or sacral spinal roots and the pelvic ganglia/pelvic nerve (Langley and Anderson, 1895; Drizenko et al., 2003; Mauroy et al., 2003; Alsaid et al., 2011; Bertrand et al., 2016). Our result is also consistent with studies in female rats, where transection of the motor branch of the pudendal nerve innervating the urethral rhabdosphincter failed to abolish the vaginocervical reflex, while a proportion of male rats recovered urinary continence after 10 days (Juárez et al., 2012).

To demonstrate how modern 3D neuroanatomy can advance spinal cord mapping, we produced a series of virtual transverse slices (six per lumbosacral segment), showing immunofluorescence for ChAT and the cytoarchitectural marker, NeuN. This revealed a limitation of the current atlases that select a single location to represent each segment (Watson et al., 2009, 2021). Specifically, the IML extends from mid-L6 to caudal S1, whereas in the atlases it is absent from L6 but present in S1 and S2. This anomaly could be caused by strain differences reported for Sprague-Dawley and Wistar rats (Pascual et al., 1989), but the section location chosen for the atlases could also have contributed, as this corresponds to a virtual slice from the rostral end of L6 that does not contain IML.

The recent atlases have revised the ontological nomenclature to identify SMN regions. Previously collectively identified as Rexed’s lamina IX (Rexed, 1952, 1954; Molander et al., 1984), subdivisions of lamina IX are now named after the dominant MN pool in the corresponding location. SMN pools in the pelvic motor system have been identified by neural tracing and functional experiments (Schrøder, 1980; Thor and de Groat, 2010). Reviewing these reports identified an anomaly in the rodent atlases, as the ExU9 nucleus that targets the urethral rhabdosphincter (and ischiocavernosus muscle in males) is shown in the ventromedial column and not the ventrolateral column, the latter being the correct location supported by strong functional anatomical evidence from multiple studies (Schrøder, 1980; Vizzard et al., 1995; Nadelhaft and Vera, 1996, 2001). The revised nomenclature is simple and clear but can be misleading, as other SMN pools can be colocated, as is the case of pelvic SMNs where motor pools are partly split across regions in different columns (Schrøder, 1980).

Spinal cord MNs are typically multipolar with long dendrites extending through a large volume of spinal cord grey and white matter (Vrieseling and Arber, 2006; Fukuda et al., 2020). In most CNS neurons, dendrites receive significantly greater input than the soma, so the perinuclear area defined by dendritic fields is important for understanding the functional anatomy of neuronal inputs. We used 3D microscopy to visualize and measure the perinuclear volume of the L6-S1 IML and found it was larger in males than females. This morphological sex difference adds to the extensive evidence of sexual dimorphism in the pelvic motor system (Jordan et al., 1982; Anderson et al., 2009; Forger, 2009; Thor and de Groat, 2010; Oti and Sakamoto, 2023).

A prominent dendritic bundle connecting the L6-S1 IML and ExU9 was identified by 3D microscopy. Dendritic bundles develop postnatally (Bellinger and Anderson, 1987; Markham et al., 1991), and in rat the largest are formed by SMNs in the lumbosacral spinal cord, with the ventrolateral bundle containing 1200-1600 dendrites (Roney et al., 1979). VMNs in the L1-L2 and L6-S1 IML also form prominent long dendritic bundles that extend medially into the DCN/SDCom and are evenly spaced to appear ladder-like in horizontal sections (Peddie and Keast, 2011). The limitation of sections has biased neuroanatomical descriptions of dendritic bundles in the lumbosacral spinal cord as they have been identified only from transverse or horizontal sections (Roney et al., 1979). 3D microscopy revealed CTB^+^ dendrites of L6-S1 VMNs that were oriented rostrodorsally and joined dendritic bundles formed by the ExU9 SMNs. A major output of the sacral IML is bladder contraction that during micturition is coordinated with inhibition of the ExU9 SMNs to relax the urethral rhabdosphincter. Therefore, the colocation of autonomic and somatic dendrites in bundles could have functional significance. For example, there is evidence suggesting electrical coupling of bundled dendrites can synchronize SMN activity and their output to functionally coupled groups of striated muscles (Roney et al., 1979; Personius et al., 2007).

We focused on the pelvic motor system in rat lumbosacral spinal and could only partly explore the neuroanatomical detail available in these rich datasets. This was highlighted by the resolution of the raw 3D images revealing single axons projecting from sacral SMNs in the margin of the ventrolateral gray matter that entered the white matter and then abruptly turned and ran for long distances in the white matter tracts. This could correspond to sacral motor neuron pools that have ascending intersegmental inputs to drive activity in lumbar locomotor pattern generators (Cazalets and Bertrand, 2000; Sourioux et al., 2018; Mille et al., 2021).

## Acknowledgements

We acknowledge the Biological Optical Microscopy Platform (BOMP), University of Melbourne (RRID:SCR_018888), for providing training and access to the light sheet microscope, and the Melbourne Histology Platform for access to cryostats. Luke Bowden performed the counts of preganglionic neurons for the CTB experiments. Research reported in this publication was supported by the Office of the Director, Stimulating Peripheral Activity to Relieve Conditions (SPARC) Program (sparc.science, RRID:SCR_017041), National Institutes of Health under Award Number OT2OD023872. The content is solely the responsibility of the authors and does not necessarily represent the official views of the National Institutes of Health.

